# Structure, inhibition, and regulation of two-pore channel TPC1 from *Arabidopsis thaliana*

**DOI:** 10.1101/041400

**Authors:** Alexander F. Kintzer, Robert M. Stroud

## Abstract

Two-pore channels (TPCs) comprise a subfamily (TPC1-3) of eukaryotic voltage-and ligand-gated cation channels^1,2^ that contain two non-equivalent tandem pore-forming subunits that then dimerize to form quasi-tetramers. Found in vacuolar^3^ or endolysosomal^4^ membranes, they regulate the conductance of sodium^5^ and calcium^3,6^ ions, intravesicular pH^5^, trafficking^7^ and excitability^8,9^. TPCs are activated by a decrease in transmembrane potential^1,3,9,10^, an increase in cytosolic calcium concentrations^1,10^, and inhibited by luminal low pH, and calcium^11^, and regulated by phosphorylation^12,13^,. We report the crystal structure of TPC1 from *Arabidopsis thaliana* (TPC1) at 2.8×4.0×3.3Å resolution as a basis for understanding ion permeation^3,4,10^, channel activation^1,5,10^, the location of voltage-sensing domains^1,9,10^, and regulatory ion-binding sites^11,14^. We determined sites of phosphorylation^3,4^ in the N-terminal and C-terminal domains that are positioned to allosterically modulate cytoplasmic Ca^2+^-activation. One of the two voltage sensing domains (VSDII) encodes voltage sensitivity and inhibition by lumenal Ca^2+^ and adopts a conformation distinct from the activated state observed in structures of other voltage-gated ion channels^15,16^. The structure shows that potent pharmacophore trans-NED19^17^ allosterically acts by clamping the pore domains to VSDII. In animals NED 19 prevents infection by Ebola virus and Filoviruses presumably by altering their fusion with the endolysosome, and delivery of their contents into the cytoplasm^7^ (Supplementary Discussion).

Diffraction was optimized and the final conditions depended on relipidation, partial dehydration, NED19^17^, and deletion of residues 2-11. The structure was determined *de novo* to 2.87Å by 9 metal substitutions and derivatives, and refined to R_cryst_=29.7%, R_free_= 33.9% (Methods and Extended Data Table 1,2,3).

TPC1 consists of two non-identical Shaker-like^18,19^ pore-forming subunits, domain I and II, separated by an EF-hand domain (Fig. 1). The central pore domains PI (S5-S6) and PII (S11-S12) couple the voltage-sensing domains VSDI (S1-S4) and VSDII (S7-S10) and modulatory cytosolic N-terminal (NTD), the central EF-hand (EF), and C-terminal (CTD) domains, which extend 20Å into the cytoplasm (Fig. 2, Extended Data Fig. 1, Methods for detailed domain assignments). The experimentally determined electron density unbiased by any model (Extended Data Fig. 2), showed that TPC1 forms a rectangular structure, ~100Å × 70Å in the membrane plane (Fig. 1). During review of our manuscript and after referee reviews, we became aware or the preceeding manuscript that reports the crystal structure of the same molecule, accompanied by extensive functional characterizations^10^. The structures are generally in excellent agreement. We additionally observe the CTD, a lipid, the inhibitor NED19, and there is some variation in observed ion binding sites. We discuss our results in light of their observations.

**Figure 1.**
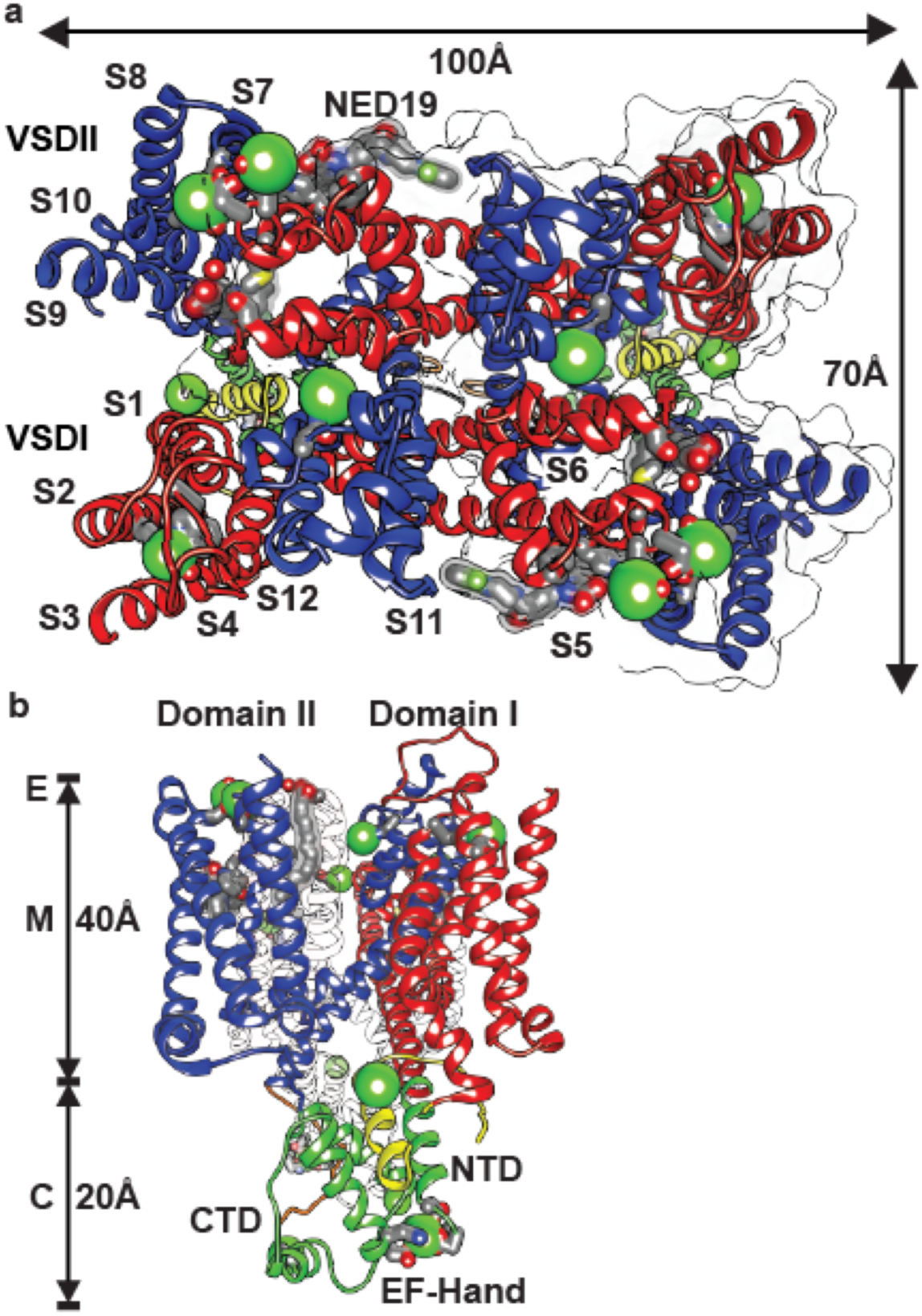
Overview of the TPC1 Structure. **a**, Top view from the lumenal side onto the membrane plane and **b**, side view from the right side with the perpendicular to the membrane plane vertical of TPC1 structure labeled by domain and transmembrane helices. E, M, and C denote approximate endolysosomal, membrane, and cytosolic boundaries. The positions of Ca ion binding sites, bound lipids, and NED 19 are shown. Approximate geometric dimensions of TPC1 are indicated.

**Figure 2.**
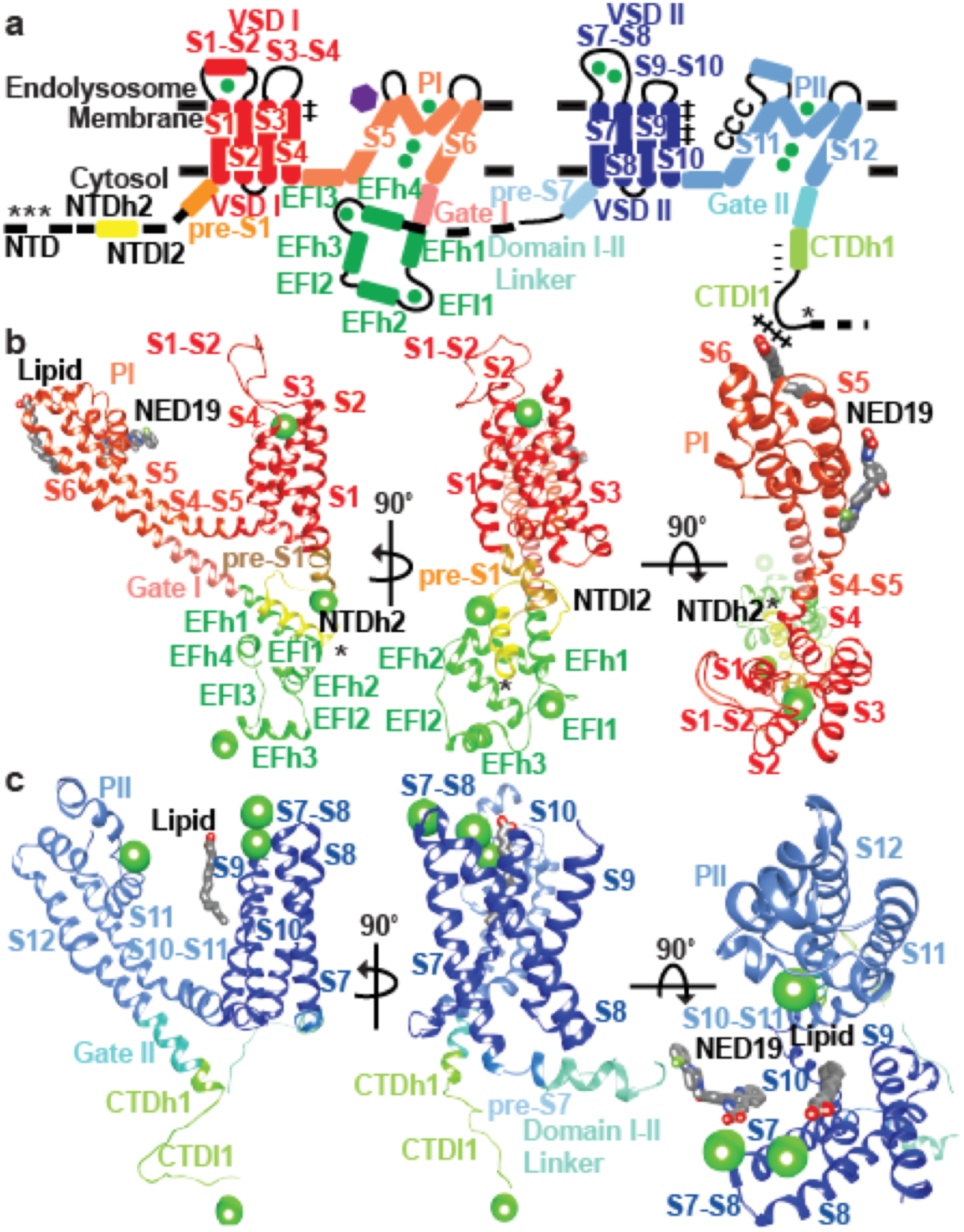
Structural details of the TPC1 monomer. Structure and domain boundaries in TPC1. **a**, A diagram describing the arrangement of structural domains, colored to match the structures below. Helices are depicted as cylinders and loops as lines. Structural details of TPC1 **b**, domain I and **c**, domain II from three perpendicular views. Phosphorylation sites are shown (*). Charges in S4 (++), S10 (++++), poly-E (----) in CTDh1, poly-R (++++) in CTDl1, and CxxCxxC (CCC) are shown. Ca^2+^ ions are shown in green and NED 19 binding site shown as a purple hexagon.

Plant and human TPCs respond to phosphoregulation by specific kinases^3,4^. We determined sites of phosphorylation in TPC1 using a combination of mass spectrometry and binding of ProDiamond-Q to N-and C-terminal truncations (Extended Data Fig. 3 and Supplementary Discussion). Residues S22, T26, and T29 in the NTD are sites of phosphorylation in TPC1. S22 is conserved in both human TPCs, whereas T26 is only found in hTPC1 (Extended Data Fig. 1). The ordered structure begins with NTD helix 2 (NTDh2; residues P30-G46). NTDh2 makes hydrophobic and polar contacts with the EF-hand domain helix 1 (EFh1), EFh2, EFh4, and VSDI (S2-S3), leading into a NTD loop 2 (NTDl2) that makes nearly a 180° turn to connect to the pre-S1 helix of VSDI. The proximity of the NTD phosphorylation sites to the Ca^2+^-activation site in the EF-hand^14,10^ EFh3 and loop EFl3, and VSDI suggests that phosphorylation in the NTD could modulate channel opening by influencing the local structure of the Ca^2+^ site or conformation of VSDI via a salt-bridge (E50-R200) between NTDl2 and VSDI (S4-S5).

Plant and human TPC1s form voltage-dependent inwardly-rectifying (towards the cytosol) ion channels, whereas hTPC2 is voltage insensitive^1,5,9,10,12^. In contrast to Ca_v_, Na_v_ K_v_ channels, activation of TPC channels is characteristically much slower, and they do not inactivate hence the original naming for plant TPCs, as slow vacuolar channels^1,5,9,10,12^.

Voltage-sensing is primarily encoded by VSDII and not by VSDI (Fig. 3)^10^. VSDII contains a classic voltage-sensing motif with four arginines R531, R537, R540, R543 in S10 (Fig. 3b, Extended Data Fig. 4a) and a conserved counter anion charge-transfer center in (Y475-E478-R537) in S8 (Extended Data Fig. 4b), as in NavAb^16^. Three other arginines form corresponding ion pairs with anions on S9 (E511-R531, D500-R540, E494-R543). The accompanying paper shows that the three arginines R537, R540, R543 are required for voltage-sensing in TPC1^10^. The second’voltage-sensing’ arginine R537 interacts with the conserved lumenal charge transfer center near Y475^10,16^, creating an opening in the cytosolic side of VSDII, and exposing R543 to the cytosol. VSDII is in a novel conformation with respect to other voltage-gated ion channel structures, which each place the last charge, equivalent to R543, at the charge transfer center, thought to represent activated VSDs^15,16^. The lumenal/extracellular side of activated VSDs adopts a more open conformation, while the cytosolic side is narrow and closed. This may mean that the TPC1 VSDII structure represents an inactive state. VSDII is indicated as the functional voltage sensor in human TPC1 also, since the R539I mutation (corresponding to R537 in plant TPC1 VSDII) eliminates most of the observed voltage-sensitivity^9^. In the human TPC2 channel the corresponding residue is I551, consistent with its lack of voltage-sensitivity.

VSDI is structurally divergent from VSDII, containing only two arginines R185 and R191 in S4, and has no conserved charge transfer center in S2 (Fig. 3a, Extended Data Fig. 4a,b) therefore not predicted to form a functional voltage sensor. These predictions are corroborated by site-directed mutagenesis and electrophysiology^10^. Intervening lateral amphipathic helices on the cytoplasmic leaflet S4-S5 and S10-S11 link VSDI and II to the pore domains PI and PII, respectively, making polar and hydrophobic contacts with the pore gates I (Gate I) and II (Gate II), and providing connections for coupling voltage-sensing, phosphorylation, and binding of ions, inhibitors, or proteins to the pore. S4-S5 is critical for coupling voltage to the pore gates^9^ in human TPC1.

**Figure 3.**
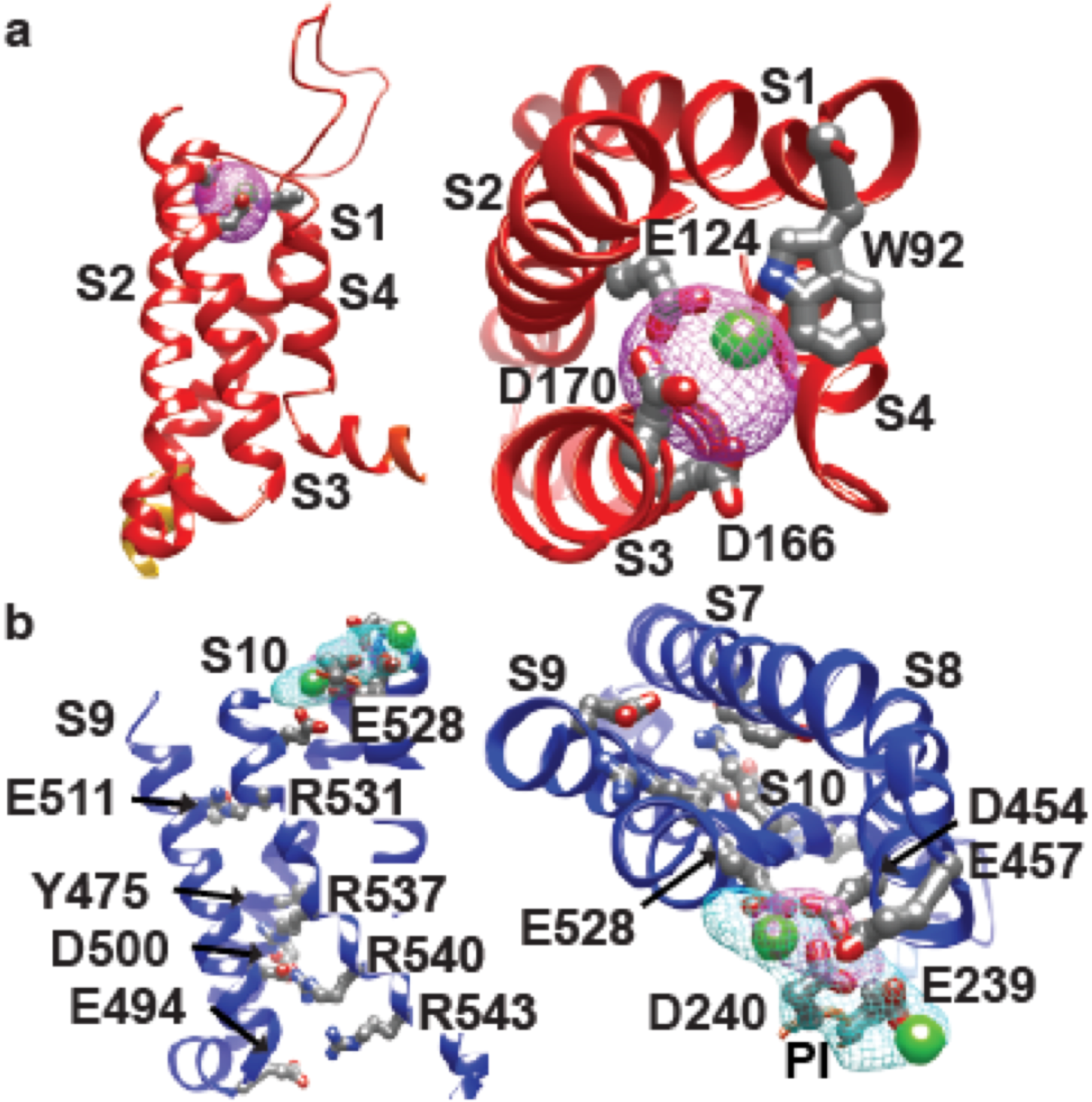
Voltage-dependence and its modulation by ions. Side (left) and top (right) views of Ca^2+^-binding sites in **a**, VSDI and **b**, VSDII. Ca^2+^ ions are shown in green. Ion coordinating residues are labeled in VSDI and II. Voltage-sensing residues are shown in VSDII. Ba^2+^ (cyan) and Yb^3+^ (magenta) isomorphous difference density peaks and atom positions contoured at 10 σ numbered according to Extended Data Fig. 2c.

Structural divergence between VSDI and II generates the rectangular shape of TPC1. Variations in the S4 angle are found in symmetrical tetrameric ion channels, though the functional relevance is not clear (Extended Data Fig. 5). In the structure of Ca_v_1.1 asymmetry of the pore domains PI-PIV plays a role in ion permeation and selectivity while the VSDs remain close to symmetrical^20^. TPCs and other tandem channels may use structural asymmetry to expand the repertoire of sensory responses and facilitate channel gating, as in the regulation of Ca_v_ channels by ions, toxins, pharmacophores, phosphorylation, or additional subunits^21^.

Lumenal Ca^2+^ inhibits plant TPC1 conductance^10,11^. The plant TPC1 mutant D454N/*fou2* in the S7-S8 loop abolishes lumenal Ca^2+^ sensitivity and increases the rate of channel opening^11^.

In our structure there are two Ca^2+^ binding sites on the lumenal side of VSDII. Site I is coordinated by E457 from S7-S8 loop and E239 of PI, whereas Site II uses D454, D240 of PI, and E528 of S10. Based on mutagenesis and patch-clamp electrophysiology of TPC1 only Site 2 is critical for lumenal inhibition by Ca^2+^ ions^10^. Heavy atom mimetics of Ca^2+^ (Ba^2+^ and Yb^3+^) validate the placement of Ca^2+^ in Sites I and II, defined by isomorphous and anomalous diffraction difference maps (Fig. 3, Extended Data Fig. 2c). A single fully occupied Yb^3+^ ion (Yb2) and Ba^2^ site (Ba1) overlaps with the Site 2. In hTPC2, E528 is an aspartate, suggesting that there may be mechanistic similarities in ion and pH sensitivities between hTPC2 and plant TPC1. VSDI binds a structural Ca^2+^ ion, confirmed by replacement with a Yb^3+^ ion (Yb1) coordinated by acidic residues E124 in S2 and D166 and D170 in S3 (Fig. 3a). Human and plant TPCs have an acidic residue E124 in S2 suggesting that this region could be a conserved Ca^2+^-binding site. However, this ion binding site has not been studied functionally.

NED19^7,17^ and Ca_v_ channel agonists and antagonists inhibit opening of TPC channels. Mutagenesis in human TPC1 demonstrated that L273 is important for channel activation and Ca^2+^-release^22^. The equivalent residues in TPC1 (M258 and L623) line PI and PII and form a hydrophobic surface that could bind hydrophobic ligands or stabilize the closed state in the absence of agonist (Extended Data Fig. 6). We observe density in TPC1 corresponding to the polycyclic TPC inhibitor NED19 (Fig. 4). Interactions with NED19 involve F229 in S5, both hydrophobic pi-stacking and a hydrogen bond between the carboxylate of NED19 and the sidechain amino group of W232 in S5, L255 in PI, F444 in S10, and W647 in S12. NED19 effectively clamps the pore domains and VSDII together, allosterically blocking channel activation. The binding site is consistent with NED19 as a high-affinity TPC-specific inhibitor^7,17^. TPC1 S6 and S12 are 25-50% conserved with S6 helices in Ca_v_ channels (Extended Data Fig. 4c). Dihydropyridines (DHP) and phenyl alkyl amines (PPA) pharmacophores bind to specific aromatic residues at a nearby, yet distinct, region in S6 of the third and fourth tandem pore forming domains in Ca_v_ channels^23^. Homologous DHP-binding residues Y288, Y656, L663, L664 in TPC1 and conserved residues in hTPC1 and hTPC2 could bind Ca_v_ pharmacophores in a parallel mechanism of inhibition^7^.

**Figure 4.**
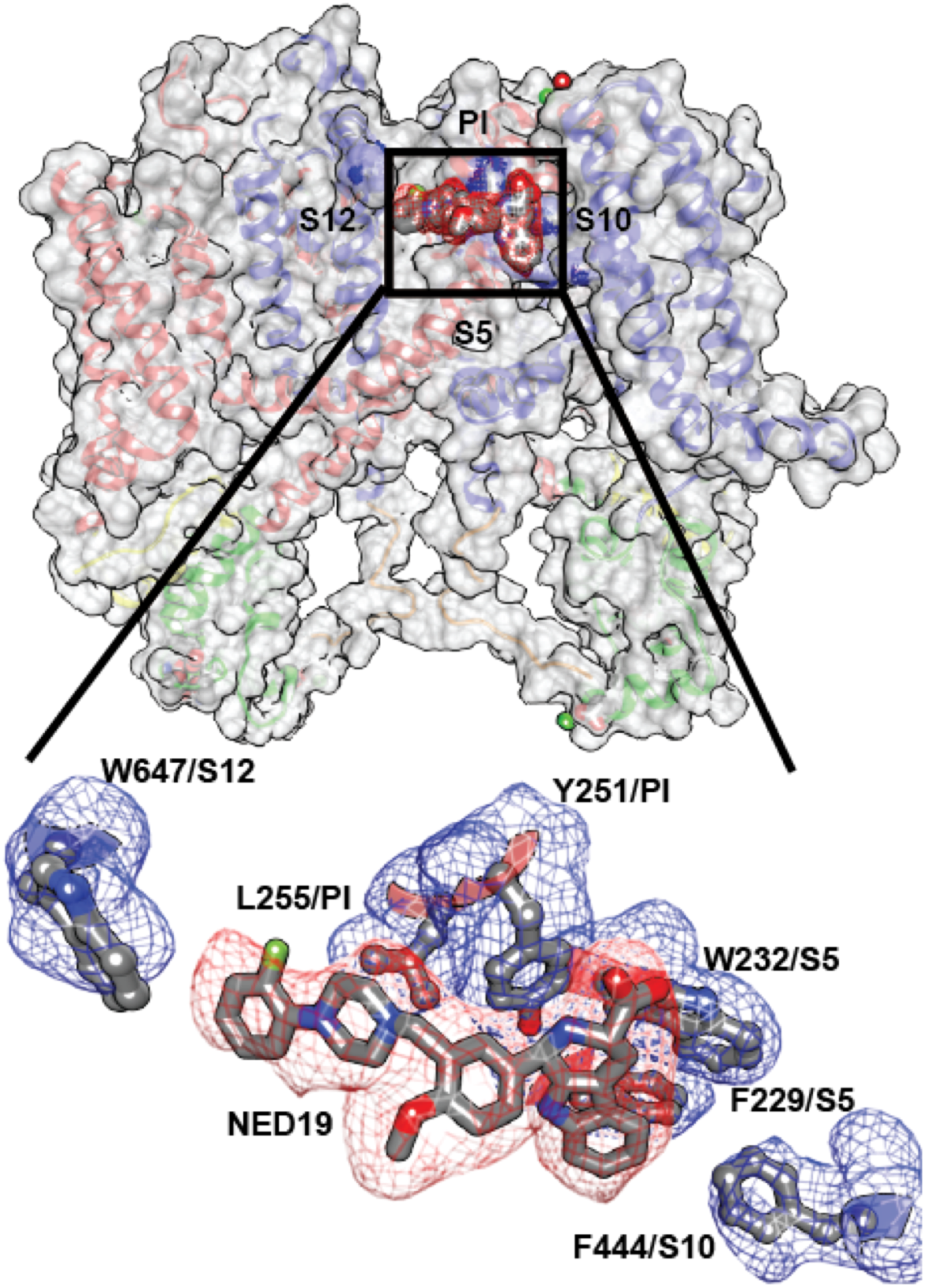
Binding site for trans-NED19. Structural interactions between the TPC1 and NED 19. (top) Surface rendering showing the location of the NED 19 binding site. (bottom) Molecular details of the NED 19 binding-pocket at the interface between PI, PII, and VSDII. Simulated-annealing omit density contoured at 1σ is shown for NED 19 (red) and interacting side chains (blue).

Ion selectivity varies among TPC channels. Plant TPCs are non-selective^3,10^, whereas human TPC1 is Na^+^-selective^5,9^, and TPC2 conducts Na^+^, Ca^2+^, and possibly H^+^ ions^5,9,24^. While the selectivity filter in TPCs is fairly conserved with Na_v_, and Ca_v_ channels, the specific anionic residues known to impart Na^+^ or Ca^2+^ selectivity^16,25^ are replaced by hydrophobic side chains (Extended Data Fig. 4d). The largely hydrophobic selectivity filter of TPC1 agrees with the reported lack of ion selectivity^10^. A single anion in the selectivity filter of hTPC1 pore loop II E642 could confer Na^+^-selectivity as this residue, N624 faces the pore lumen in TPC1. Pore loop I forms a re-entrant helix-turn-helix motif that forms a wide lining of two sides of the pore mouth (Fig. 5a). The high resolution data and crystallization conditions enabled visualization of electron density for lipid molecules, one bound to the backside of PI at the interface of VSDII, flanking the lumenal Ca^2+^-binding Site II. Modeled as a palmitic acid, the lipid is a hydrogen bond acceptor from T241 and M237 of PI and bound adjacent to D240 the site for lumenal Ca^2+^-inhibition. This could provide a basis for the reported modulation of TPCs by fatty acids. (Supplementary Discussion). By contrast, pore loop II forms an extensive asymmetric constriction site at the pore mouth that brings four negative charges D605, D606 to its lining of which D605 likely coordinates a Ca^2+^ ion (Fig. 5b), confirmed by Yb^3+^ (Yb3) observed in isomorphous and anomalous diffraction difference maps. The coordination distance of 2.5Å suggests that this site could recognize ions from the lumen. However, this site is unlikely to function in selecting ions, as the wide pore mouth ~12-15Å of PI does not provide a barrier to ion flow (Fig. 5c). D269 of PI faces the pore, but the distance to the lumen is much to far 12.6Å to function in ion selectivity.

**Figure 5.**
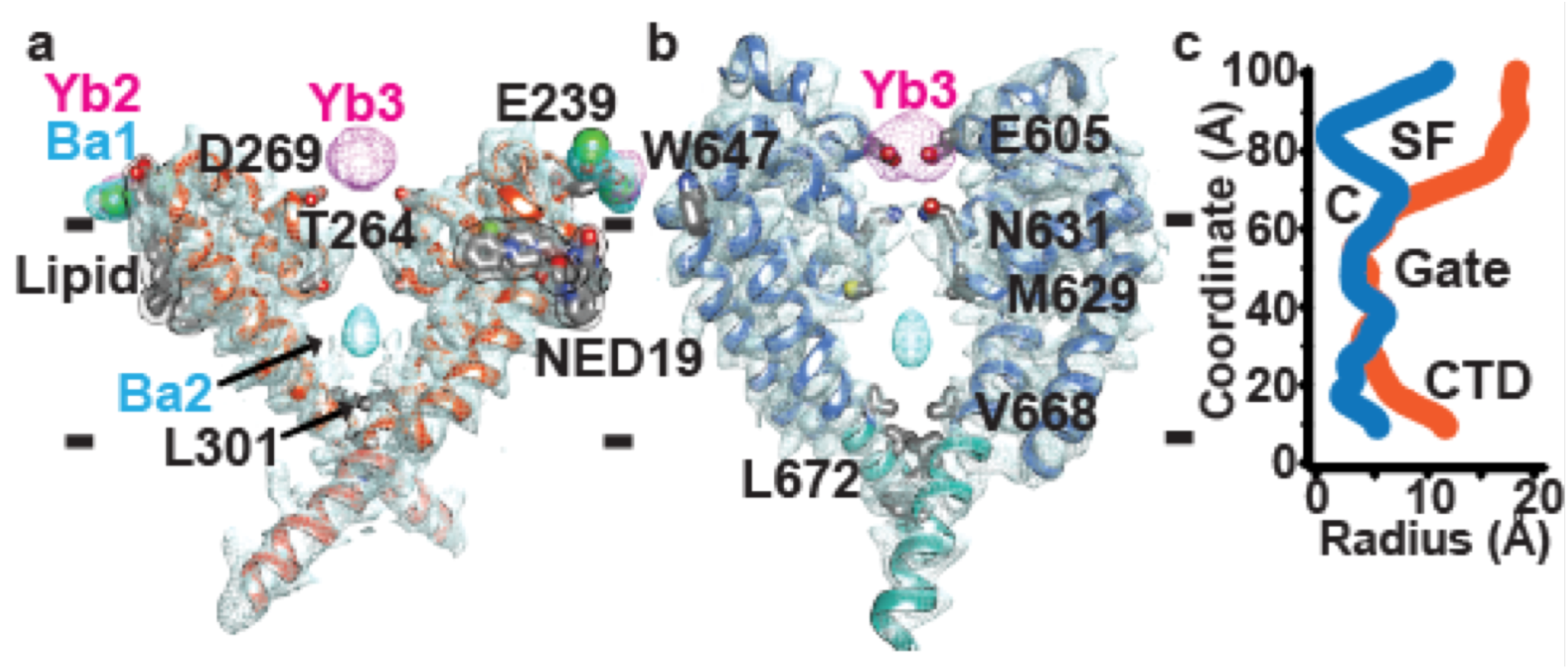
The TPC1 ion channel. Cross-sectional views of the TPC1 ion channel separately through PI and PII. Domains removed for clarity. **a**, PI to PI (orange) and **b**, PII to PII (blue) with sharpened 2mF_o_-DF_c_ electron density contoured at 1 σ and Ba^2+^ (cyan) and Yb^3+^ (magenta) isomorphous difference density contoured at 10 σ. Membrane boundaries are indicated by dashes. **c**, Pore radius calculations through separate pore domains, PI (orange) and PII (blue) using HOLE (See Methods). Approximate boundaries for the putative selectivity filter (SF), cavity (C), Gates I and II, and CTD are shown. Channel axis is vertical.

Two N631 residues converge to 4.9Å to form a narrowed region that separates the upper vestibule from the putative selectivity filter. However, opposing Q633 residues from PI are 12.2Å apart leave the gasket open to ion flow. This is the narrowest part of the selectivity filter.

Two T264 residues, conserved in Ca_v_, Na_v_, and K_v_ subunits, define the upper boundary of the central cavity in TPC1 (Extended Data Fig. 4d, Fig. 5c). The central cavity contains solvent molecules and barium ions (Ba2), defined by isomorphous and anomalous difference maps. However, no ions are observed in the selectivity filter, unlike highly selective Na_v_, Ca_v_, or K_v_ channels^16,20,15^. The central cavity is water filled and lined by hydrophobic residues. Two rings of hydrophobic residues L301, V668, Y305, and L672 seal the cavity toward the cytoplasm, and are part of the extensively studied pore gates of K_v_ channels^26^^−^^28^ and Gates I and II in TPC1. On the cytosolic face of the gates, a salt-bridge K309-E673 staples Gate I and II together. K^+^ ions have been observed in the gate and lower selectivity filter of MlotiK1^29^ and KcsA^30^ and mutation of similarly positioned hydrophobic residues in the gate can modulate the channel state or flux of ions through the channel. This supports the idea that the VSDs, selectivity filter, and gate undergo coordinated movements during channel opening and closing to regulate ion conductance.

In plant TPC1, two EF-hand domains between domains I and II, confer sensitivity to cytosolic Ca^2+^ ions^14^. These domains consist of two Ca^2+^ binding motifs EFl1 and EFl3, with 25-30% sequence conservation to human calmodulin (hCAM, Extended Data Fig. 4e). EFh1 forms a 70Å long helix continuous with the gate helix S6 (Fig. 2b, Fig. 6a) that is connected to the first Ca^2+^-binding site in EFl1 formed by sidechain interactions with D335 (2.6Å), D337 (2.8Å), N339 (3.2Å), and the backbone of E341 (2.3Å). This site is probably structural, as it does not function in Ca^2+^-activation^10,14^. The second site in EFl3 has been shown to facilitate channel activation by cytosolic Ca^2+^, as a D376A mutant abolished all Ca^2+^-dependent activation^14^. We observe an Ca^2+^ ion in EFl3 coordinated by E374 (2.9Å) and D377 (5.2Å), but surprisingly not by D376 which is 8.5Å away. Thus the mechanism of Ca^2+^-activation may involve a structural change in EFl3 involving the Ca^2+^-binding site and D376.

**Figure 6.**
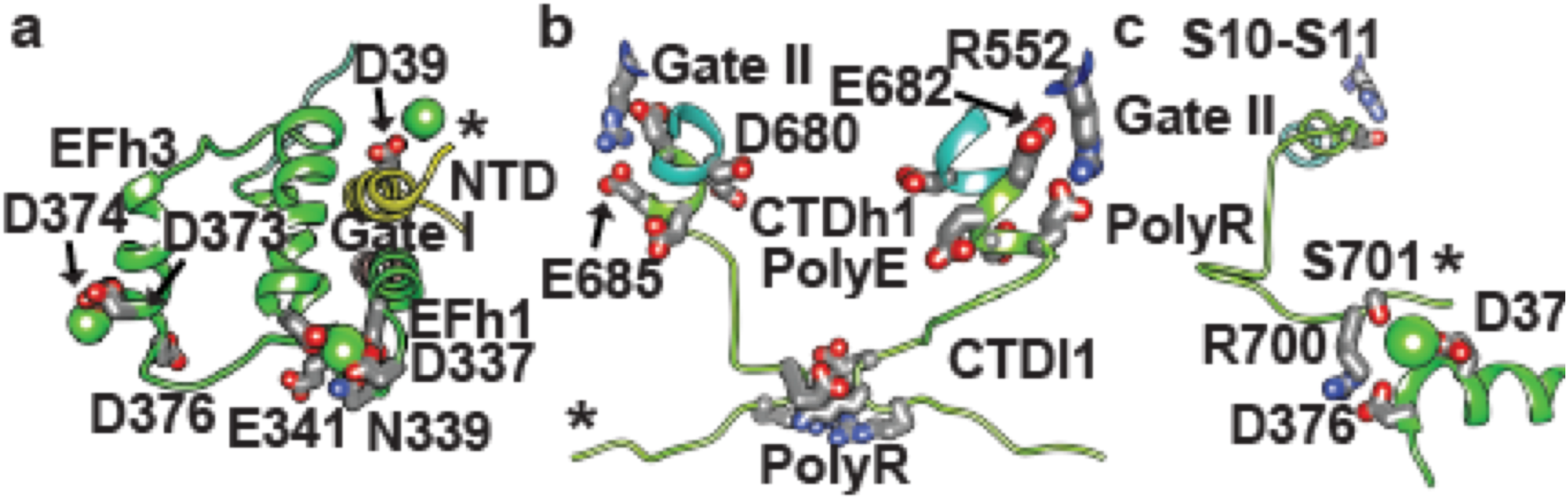
Cytosolic sensory domains. Structure of EF-hand and coupling with NTD, VSDII, and CTD. Sideviews through **a**, the EF-hand (green), CTD (chartreuse), and NTD (yellow), **b**, constriction site formed by Gate II and CTD, and **c**, interactions between EF-hand and CTD. Phosphorylation sites are indicated by stars (*). Transmembrane and adjacent domains removed for clarity.

The CTD plays a critical role in plant TPCs (Supplementary Discussion), though least conserved across species (Extended Data Fig. 1). CTD helix 1 (CTDh1) continues from Gate II and S12 with a poly-anionic (poly-E) helix E682-E685 (Fig. 6b). Ca_v_ channels contain a similar conserved poly-anionic region of variable length (Extended Data Fig. 4c). Poly-E residues E682 and E685 hydrogen bond to R552 of S10-S11, directly linking the conformation of the CTDh1 to the voltage-sensor VSDII S10-S11. CTD loop 1 (CTDl1) from each monomer converges to form a charged constriction site D694, K695, and R696 (Fig. 6b, Fig. 5c), that bring six charged sidechains to stabilize the channel through inter-monomer contacts. The intermolecular constriction site partially includes a poly-R motif formed by R696, R698, R699, R700, and may play a structural role or regulate binding of second messenger lipids or proteins (Extended Data Fig. 7). Importantly, the CTD forms an intra-molecular complex with the Ca^2+^-activation site EFl3 (Fig. 6c). Surprising, R700 from CTDl1 makes a salt-bridge with the critical residue for Ca^2+^-activation R700-D376. S701 coordinates the bound Ca^2+^ (4.2Å) in EFl3 along with E374 and D377. These interactions include the potential site of phosphorylation S706 in CTDl1, not observed in the structure. Thus, reversible phosphorylation in the CTD could disrupt the EFh3-CTDl1 interaction and modulate Ca^2+^-sensing or couple directly to the VSDII via the R552-E682-E685 interaction (Fig. 6c). The interactions between the regulatory sites and the central channel are schematized in Extended Data Fig. 7. The channel conductance of human and plant TPCs is multi-modal^3^. Our structure suggests a mechanism for channel opening, whereby Ca^2+^ concentrations, voltage, and phosphoregulation are integrated through conformational changes in the VSDs, selectivity filter, gate, NTDs, and CTDs to govern the conduction of ions (Extended Data Fig. 7).

## Methods

### Protein Production

AtTPC1 was expressed in codon-optimized form in *Saccharomyces cerevisiae* based on p423_GAL1^31^ (Uniprot accession number Q94KI8). A C-terminal Thrombin cleavage site was followed by a 12 residue glycine-serine linker and a 10 residue histidine tag. Transformed S. cerevisiae (strain DSY-5) were grown from overnight starter cultures in synthetic complete defined medium lacking histidine (CSM-HIS) in 1L shaker flasks at 30°C to OD_600nm_ of 12-15. Protein expression was induced by adding 8% Galactose dissolved in 4x YP medium, to a final concentration of 2% Galactose, at 30°C for 18-20hr. Cells were harvested by centrifugation, resuspended in lysis buffer (50 mM Tris pH 7.4, 500 mM NaCl, 1 mM EDTA, 1 mM PMSF), followed by bead beating using 0.5mm beads for 30 minutes at 1 minute intervals. The lysate was then centrifuged at 21,000g for 20 minutes, followed by collection of membranes by centrifugation at 186,010g for 150 minutes. Membranes were resuspended (50 mM Tris pH 7.4, 500 mM NaCl, 10% Glycerol), aliquoted, and frozen in liquid nitrogen for storage at −80°C. 7-12 g aliquots of membranes were solubilized in membrane buffer containing 1% (0.2 g/g of membrane) n-dodecyl-beta-D-maltopyranoside (DDM), 0.1 mM cholesteryl hemisuccinate (CHS), 0.03 mg/mL soy polar lipids, 1 mM CaCl_2_, and 1 mM PMSF, for 1.5 hours at 4°C. The solubilized material was collected by centrifugation at 104,630g for 20 minutes, filtered through a 5μm filter, and incubated with 3 mL of NiNTA beads in the presence of 20 mM Imidazole pH 7.4 for 7.5 hours at 4°C. NiNTA beads were collected by draining the mixture through a column and the beads were washed successively with 25 mL of NiNTA wash buffer (50 mM Tris pH 7.4, 500 mM NaCl, 5% Glycerol, 0.1% DDM, 0.1 mM CHS, 0.03 mg/mL soy polar lipids, 1 mM CaCl_2_) plus 20 mM Imidazole pH 7.4 for 20 minutes at 4°C, then 25 mL of NiNTA wash buffer plus 75 mM Imidazole pH 7.4 for 10 minutes at 4°C, followed by 10 mL of NiNTA elution buffer (50 mM Tris pH 7.4, 200 mM NaCl, 5% Glycerol, 0.1% DDM, 0.1 mM CHS, 0.03 mg/mL soy polar lipids, 2 mM CaCl_2_). TPC1 protein was obtained by incubating NiNTA beads with 200 units of bovine thrombin in 25 mL of NiNTA elution buffer overnight at 4°C. The following day, NiNTA beads were washed with an additional 25 mL of NiNTA wash buffer plus 20 mM Imidazole pH 7.4 for 20 minutes at 4°C, concentrated, and loaded on a Superose 6 column equilibrated in Size exclusion buffer (20 mM Hepes pH 7.3, 200 mM NaCl, 5% Glycerol, 0.05% DDM, 0.1 mM CHS, 0.03 mg/mL soy polar lipids, and 1 mM CaCl_2_). Peaks fractions were pooled and concentrated to 5 mg/mL for crystallization. TPC1 and mutants were prepared similarly, except for the S701C mutant where 1mM and 100μM TCEP were included during solubilization and final size exclusion chromatography, respectively.

### Crystallization

Several eukaryotic TPCs, including human hTPC1, hTPC2, and AtTPC1 (TPC1) were evaluated for expression in *S. cerevisiae* and SF9 cells, and homogeneity by fluorescence-based and conventional size-exclusion chromatography. TPC1 was selected based on expression level, stability, homogeneity and crystallization trials. Nevertheless, detergent-solubilized TPC1 was unstable and resistant to crystallization. Thermostability screening^32^ identified cholesteryl hemisuccinate (CHS), lipids, and Ca^2+^ ions as improving TPC1 stability and homogeneity.

Wild-type TPC1 crystals were obtained in the presence of CaCl_2_, CHS, and soy polar lipids. Initial crystals from 200nL drops in sparse matrix trays diffracted to ~10Å resolution, with few diffracting to 4Å. Reproducibility in 2μL drops was achieved through extensive screening trials. To improve diffraction, truncation constructs were made. A construct lacking the N-terminal residues 2-11 (TPC1) diffracted on average to 7Å, with 1% showing diffraction to 3.9Å resolution, but were too radiation sensitive and anisotropic to collect complete datasets. The TPC inhibitor trans-NED19 (NED 19) was added to 1mM final, 1 hour before crystallization. These diffracted to ~5Å resolution, with 50% to 3.9Å resolution. Adding 100μL of protein and 1mg of DDM to a thin-film of 0.03 mg soy polar lipids with stirring overnight at 4°C led to the TPC1 high lipid-detergent (HiLiDe)^33^ mixture following ultracentrifugation for 20 minutes in a TLA-55 rotor at 112,000g. Average diffraction was 4Å, with 10% diffracting to 3.5-3.7Å resolution, the best being 3.5 × 5.5 × 4.0Å. Anisotropic resolution was determined using the CCP4 program Truncate^34^ and the UCLA Anisotropy Server^35^. Large improvements were achieved through partial dehydration. HiLiDe crystals were incubated with additional amounts of polyethylene glycol 300, 0-20 % (v/v), prior to harvesting. Incubating with additional 15% PEG300, increased the average diffraction to 4Å, best diffraction to 2.7Å resolution, and diminished the diffraction anisotropy. Derivative crystals were obtained by soaking in 0.1-50 mM of heavy-atom compounds in a 1:1 mixture of Crystal buffer (0.1M Glycine pH 9.3, 50 mM KCl, 1mM CaCl_2_, 35% PEG300) for 24-48 hours at 20°C. Partial dehydration was performed after derivatization.

### Data Collection and Reduction

X-ray diffraction was collected at the Advanced Light Source Beamlines (ALS) 8.3.1, 5.0.2 and the Stanford Synchrotron Radiation Laboratory Beamline (SSRL) 12-2. Data were reduced using XDS^36^. Native data were collected at wavelength 1.000Å. Data from heavy atom derivative crystals were collected at the anomalous peak of the L-III edge for each element (Ba=1.750Å, Hg=1.006Å, Ta=1.2546Å, Yb=1.3854Å).

### Structure Determination of TPC1

De novo structure determination was based on 9 heavy metal and cluster derivatives at 3.5Å (Extended Data Table 1). Dehydration improved diffraction to 2.7Å while diminishing the anisotropy (Extended Data Table 2, Extended Data Table 3). Experimental phases were calculated by multiple isomorphous replacement with anomalous scattering (MIRAS) from both non-dehydrated (Extended Data Table 1) and dehydrated (Extended Data Table 2) crystals derivatized with tantalum bromide clusters (Ta_6_Br_12_), barium(II) chloride, ytterbium(III) chloride, and a mutant of TPC1 derivatized with mercury(II) chloride.

A single tantalum atom was identified by Single Isomorphous Replacement (SIR) with Anomalous Scattering (SIRAS) using SHELXC/D^37^. A Ta6Br12 cluster was refined and phases calculated using SHARP^38^. These phases were used to place sites in other derivatives and calculate phases using SHARP^38^. Initial MIRAS phases were determined with non-dehydrated crystals with 100μM Ta6Br12 clusters (Derivative 1; 1 site, Isomorphous phasing power (PP_iso_; defined as PP_iso_ = RMS (| F_H_ |_calc_/ Σ || F_PH_ - F_P_ |_obs_ - | F_PH_ - F_P_ |_calc_|) or RMS (| F_H_ |_calc_/ Σε_iso_), where |F_PH_| is the heavy atom structure factor amplitude, | F_PH_ | the heavy-atom derivative structure factor amplitude, | F_P_ | the native structure factor amplitude, and | F_H_ | is the heavy atom structure factor amplitude, and ε_iso_ the phase-integrated lack of closure for isomorphous differences) of 1.0/0.77 (ancentric/centric); Anomalous phasing power (PP_ano_; defined as PP_ano_ =RMS (Σ |F_PH+_ - F_PH-_|_calc_/ Σε_ano_) or RMS (Σ | F_PH+_ - F_PH-_ |_calc_/ Σ||F_PH+_ - F_PH-_|_obs_ - | F_PH_+ - F_PH-_|_calc_|), where |F_PH+_ - F_PH-_|, is the mean anomalous difference between Friedel pairs, and s_ano_ the phase-integrated lack of closure for anomalous differences) of 0.31; Isomorphous Rcullis (Rcullis_iso_; defined as Rcullis_iso_ = Σ ε_iso_ / Σ | F_PH_ - F_P_ |obs) of 0.766/0.755 (ancentric/centric); Anomalous Rcullis (Rcullisan_o_; defined as Rcullis_ano_ = Σ ε_ano_ / Σ | F_PH+_ - F_PH-_ |obs of 0.584), 1mM Ta6Br12 clusters (Derivative 2; 1 site, PP_iso_ of 2.32/1.94 (ancentric/centric), PP_anom_ of 0.761, Rcullisiso of 0.525/0.536 (ancentric/centric), Rcullisano of 0.792), or 2mM Ta6Br12 clusters (Derivative 3; 1 site, PP_iso_ of 0.89/1.1 (ancentric/centric), PP_anom_ of 1.36, Rcullis_iso_ of 0.850/0.765(ancentric/centric), Rcullis_ano_ of 0.572), using merged data (Native 1) from three non-dehydrated crystals as a source of isomorphous native amplitudes (Extended Data Table 1).

This map allowed building of all transmembrane segments and partial building of the EF-hand domain in TPC1. Sidechain features and density for loop regions were largely absent from calculated maps, precluding subunit identification, and objective determination of the helical register. Initial MIRAS phases were further improved by including merged data from five non-dehydrated crystals derivatized with 50mM barium(II) chloride (Derivative 4; 3 sites, PP_iso_ of 0.75/0.58 (ancentric/centric), PP_anom_ of 0.59, Rcullis_iso_ of 0.905/0.888 (ancentric/centric), Rcullis_ano_ of 1.00) in the MIRAS calculation, resulting in an overall FOM of 0.262/0.333 (ancentric/centric) (Extended Data Table 1). Combined tantalum and barium phases allowed fitting of some sidechains to density, placement of ions binding sites in the channel lumen and VSDs (Extended Data Fig. 2, Supplementary Data Table 1). The best source of phases came from derivatives obtained from crystals that were dehydrated after a 24-48 hour soak in heavy atom solution. MIRAS phases were calculated from native crystals derivatized with 1mM Ta_6_Br_12_ clusters (Derivative 5; 1 sites, PP_iso_ of 4.09/3.56 (ancentric/centric), PP_anom_ of 2.42, Rcullis_iso_ of 0.302/0.286 (ancentric/centric), Rcullis_ano_ of 0.495) and Derivative 6; 1 sites, PP_iso_ of 8.54/8.91 (ancentric/centric), PP_anom_ of 2.29, Rcullis_iso_ of 0.149/0.122 (ancentric/centric), Rcullis_ano_ of 0.517) or 2mM Ta_6_Br_12_ clusters (Derivative 7; 1 sites, PP_iso_ of 3.65/3.54 (ancentric/centric), PP_anom_ of 2.25, Rcullis_iso_ of 0.372/0.323 (ancentric/centric), Rcullis_ano_ of 0.535), 1mM ytterbium(III) chloride (Derivative 8; 4 sites, PP_iso_ of 0.851/0.795 (ancentric/centric), PP_anom_ of 0.699, Rcullis_iso_ of 0.962/0.884 (ancentric/centric), Rcullis_ano_ of 0.982)), and a TPC1 S701C mutant derivatized with 1mM mercury(II) chloride (Derivative 9; 5 sites, PP_iso_ of 0.792/0.822 (ancentric/centric), PP_anom_ of 0.736, Rcullis_iso_ of 0.935/0.704 (ancentric/centric), Rcullis_ano_ of 1.02), followed by dehydration resulting in a FOM of 0.267/0.341 (ancentric/centric) (Extended Data Table 2). Non-isomorphism errors were minimized by using merged data from three non-dehydrated crystals extending to 3.5Å resolution (Native 1) as a source of isomorphous native amplitudes for all phasing calculations, density modification, and initial model building (Extended Data Table 1, Extended Data Table 2). MIRAS phases from dehydrated derivatives extend to higher resolution, contain less diffraction anisotropy, and display much clearer electron density for mainchain and sidechain features, and allow objective determination of helical register (Extended Data Fig. 2b). Mercury heavy atom positions were used as restraints, along with sequence homology, for objective placement of cysteine residues (Supplementary Data Table 1). Ytterbium and barium heavy atom positions were used in conjunction with homology for placing likely (D, E) chelate residues within the pore, VSDs, and pore (Extended Data Fig. 2c, Supplementary Data Table 1). MIRAS phases from non-dehydrated and dehydrated derivatives were further improved through solvent flipping in SOLOMON^39^ with 75% and 70% solvent content and FOMs of 0.637 and 0.470. Phases were then combined and extended to 3.5Å resolution through iterative cycles of density modification in DMMULTI^40^, using histogram matching, solvent flattening, gradual phase extension^41^, and 3-fold cross-crystal averaging with un-phased, dehydrated, native amplitudes (Native 2) with 70% solvent content with a FOM of 0.493 (Extended Data Fig. 2b). Averaging masks were generated by NCSMASK^34^ with radius of 4Å from atomic models from each iteration of building and refinement. Model phases were purposely left out of experimental density modification procedures to reduce bias.

Initial model building was guided by the position of heavy atoms determined from isomorphous difference maps (Extended Data Fig. 2c, Supplementary Data Table 1). Using anomalous differences yielded equivalent maps and structural restraints.

### Refinement

The best native data (Native 2) extends to overall resolution 2.8 × 4.0 × 3.3Å (ref.^35^), and enabled building of 85% of the TPC1 protein sequence. Structure interpretation was using COOT^42^, with refinement in PHENIX^43^. Simulated-annealing was employed alongside refinement of coordinates, TLS restraints, and several cycles of automated building using ARP/wARP^34^ in CCP4. Building of the NTD, CTD, and loop regions were guided by restraints based on heavy atom positions (Supplementary Data Table 1), omit maps, and gradual improvement of the map and crystallographic Rfactors. Refinement of the structure in triclinic space group P1 led to a substantial reduction in R_free_ (39.9%). During review of our manuscript and after referee reviews, we became aware or the accompanying manuscript that reports the crystal structure of the same molecule, under review at the same time, at the same journal. The authors made their coordinates of atTPC1 (PDBID 5E1J) available^10^. The topology of the two structures were identical and their analysis had determined the registration of ~14 additional sites by mutation of residues to cysteine. Therefore we checked our structure for registration against theirs. This led us to several corrections of registration in the sequence and led to improvements in map quality and crystallographic Rfactors (Extended Data Table 3). Anisotropy corrected data (2.8 × 4.0 × 3.3 Å) were used during the final stages of refinement with an applied isotropic B-factor of −78.72 Å^2^. Our structure of TPC1 was refined to 2.87Å resolution with final R_work_/R_free_ of 29.71% and 33.89%. Analysis by Molprobity shows Ramachandran geometries of 91.8%, 6.9%, and 1.3% for favored, allowed, and outliers. The structure contains 17.0% rotamer outliers.

### Domain assignment

Based on experimental electron density (Extended Data Fig. 2b), heavy atom restraints (Extended Data Fig. 2c and Supplementary Data Table 1), and sequence homology (Extended Data Fig. 1), we have assigned the TPC1 domain structure as follows (Fig. 2a, Extended Data Fig. 1): N-terminal domain (NTD; residues 30-55), pre-S1 helix (pre-S1; residues 60-71), Voltage Sensing Domain I (VSDI; S1-S4, residues 72-215), pore domain I (PI;S5-S6, residues 216-301), gate I (residues 302-321), EF-hand (EF; residues 322-399), Domain I-II linker (residues 400-418), pre-S7 helix (pre-S7; residues 419-433), VSDII (S7-S10; residues 434-564), pore domain II (PII; S11-S12, residues 565-668), gate II (residues 669-685), C-terminal domain (CTD; residues 686-703). Unstructured regions include part of the NTD (residues 1-29,56-59), S3-S4 (residues 174-183), Domain I-II linker (residues 408-412), S9-S10 (residues 519-523), PII (residues 591-593), and CTD (residues 704-733) for a total of 84 missing residues (11.5% unstructured). The structure of TPC1 comprises 649 residues (88.5% of total sequence). Supplementary Data Table 1 summarizes the restraints used for structure determination.

Sequence alignments were performed using MUSCLE^44^ and ALINE^45^. Structural figures were made using CHIMERA^46^ and PyMOL. Pore radius calculations using HOLE^47^.

## Acknowledgements

We are grateful to J. Holton and G. Meigs at ALS beamline 8.3.1 and A. Gonzalez at SSRL beamline 12-2 for help with data collection, B. P. Pedersen, B. H. Schmidt, and J. Finer-Moore for critical analysis of the manuscript. This work was supported by NIH grant GM24485 to R.M.S. Beamline 8.3.1 at the Advanced Light Source is operated by the University of California Office of the President, Multicampus Research Programs and Initiatives grant MR-15-328599 and Program for Breakthrough Biomedical Research, which is partially funded by the Sandler Foundation. The Berkeley Center for Structural Biology is supported in part by the National Institutes of Health, National Institute of General Medical Sciences, and the Howard Hughes Medical Institute. The Advanced Light Source is supported by the Director, Office of Science, Office of Basic Energy Sciences, of the U.S. Department of Energy under Contract No. DE-AC02-05CH11231. Use of the Stanford Synchrotron Radiation Lightsource, SLAC National Accelerator Laboratory, is supported by the U.S. Department of Energy, Office of Science, Office of Basic Energy Sciences under Contract No. DE-AC02-76SF00515. The SSRL Structural Molecular Biology Program is supported by the DOE Office of Biological and Environmental Research, and by the National Institutes of Health, National Institute of General Medical Sciences (including P41GM103393). The contents of this publication are solely the responsibility of the authors and do not necessarily represent the official views of NIGMS or NIH. Figures were made with Chimera developed by the Resource for Biocomputing, Visualization, and Informatics at the University of California, San Francisco (supported by NIGMS P41-GM103311). And PyMOL developed by the late Warren DeLano. Mass spectrometry was carried out by the UCSF mass spectrometry facility (supported by NIGMS P41- RR001614). The authors declare no competing financial interests.

## Author Contributions

A.F.K performed the experiments, collected and processed the data, determined, refined, and analyzed the structure. R.M.S. supervised the project, and aided in structure determination and analysis. A.F.K and R.M.S. wrote the manuscript.

## Author Information

Coordinates and structure factors have been deposited in the Protein Data Bank with the accession number 5DQQ. Correspondence and requests for materials should be addressed to R.M.S.

---

**Extended Data Figure 1.**
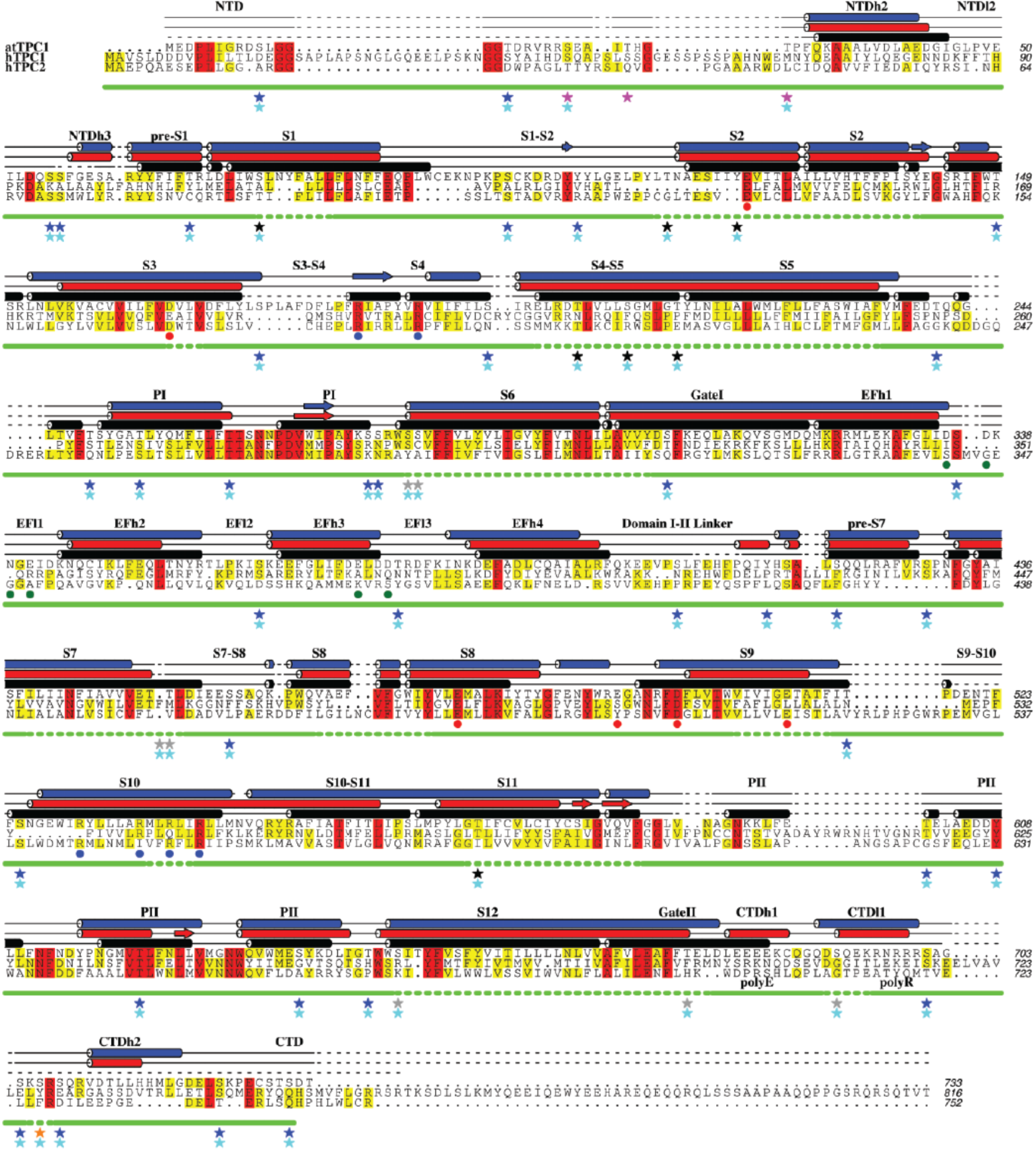
Sequence alignment of TPCs with TPC1 experimental and predicted secondary structure. A sequence alignment based on seven human and plant TPC orthologs with observed TPC1 (black) and secondary structure predictions by Psipred^48^ (red) and Jpred^49^ (blue). Helices are shown as cylinders, beta sheets as planks, coil as solid lines, and unstructured regions as dashed lines. Level of conservation is indicated by color (>50% yellow; >80% red). Blue dots (•) mark arginines in S4. Red dots (•) mark charge-transfer anions. Green dots (•) mark Ca^2+^-binding sites in the EF-hand. Solid and dashed green lines represent observed and absent peptides from mass spectrometry experiments. Magenta stars (⋆), orange stars (⋆), cyan stars (⋆), blue stars (⋆), black stars (⋆), and grey stars (⋆) mark observed phosphorylation sites using ESI-MS (Extended Data Fig. 3a), potential phosphorylation sites identified by truncation constructs (Extended Data Fig. 3e), predicted phosphorylation sites by NetPhosK^50^, non-phosphorylated sites, unlikely sites due to solvent inaccessibility, and mark unknown or unidentified regions, respectively.

**Extended Data Figure 2.**
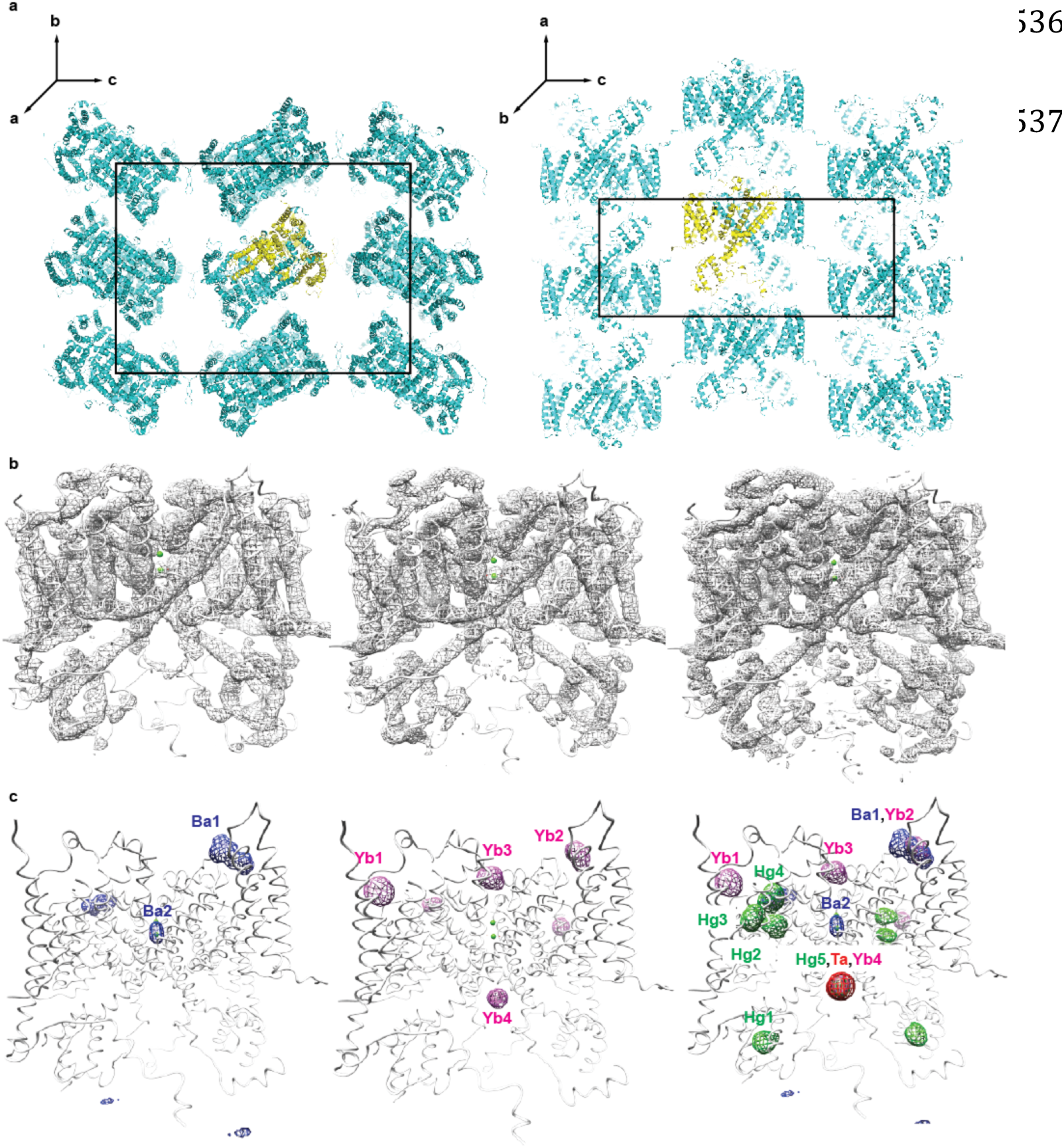
Crystal packing of TPC1 and experimental electron densities. **a**, Views of the TPC1 C2221 crystal lattice viewed along 2-fold axes parallel to **<u>a</u>** (left) and **<u>b</u>** (right). Unit cell boundaries are shown. The asymmetric unit is shown in yellow. **b**, Transverse view of TPC1 is shown with overlayed FOM-weighted experimental electron density calculated using native amplitudes (Native 1) and heavy-atom phases contoured at 1 σ. (left to right) Density modified phases from non-dehydrated derivatives (Extended Data Table 1), dehydrated derivatives (Extended Data Table 2), and combined phases with solvent flattening, histogram matching, phase extension to high resolution native (Native 2), and cross-crystal averaging (See Methods, Extended Data Table 3). **c**, Transverse views of TPC1 with overlayed heavy atom electron densities calculated from isomorphous differences.

**Extended Data Figure 3.**
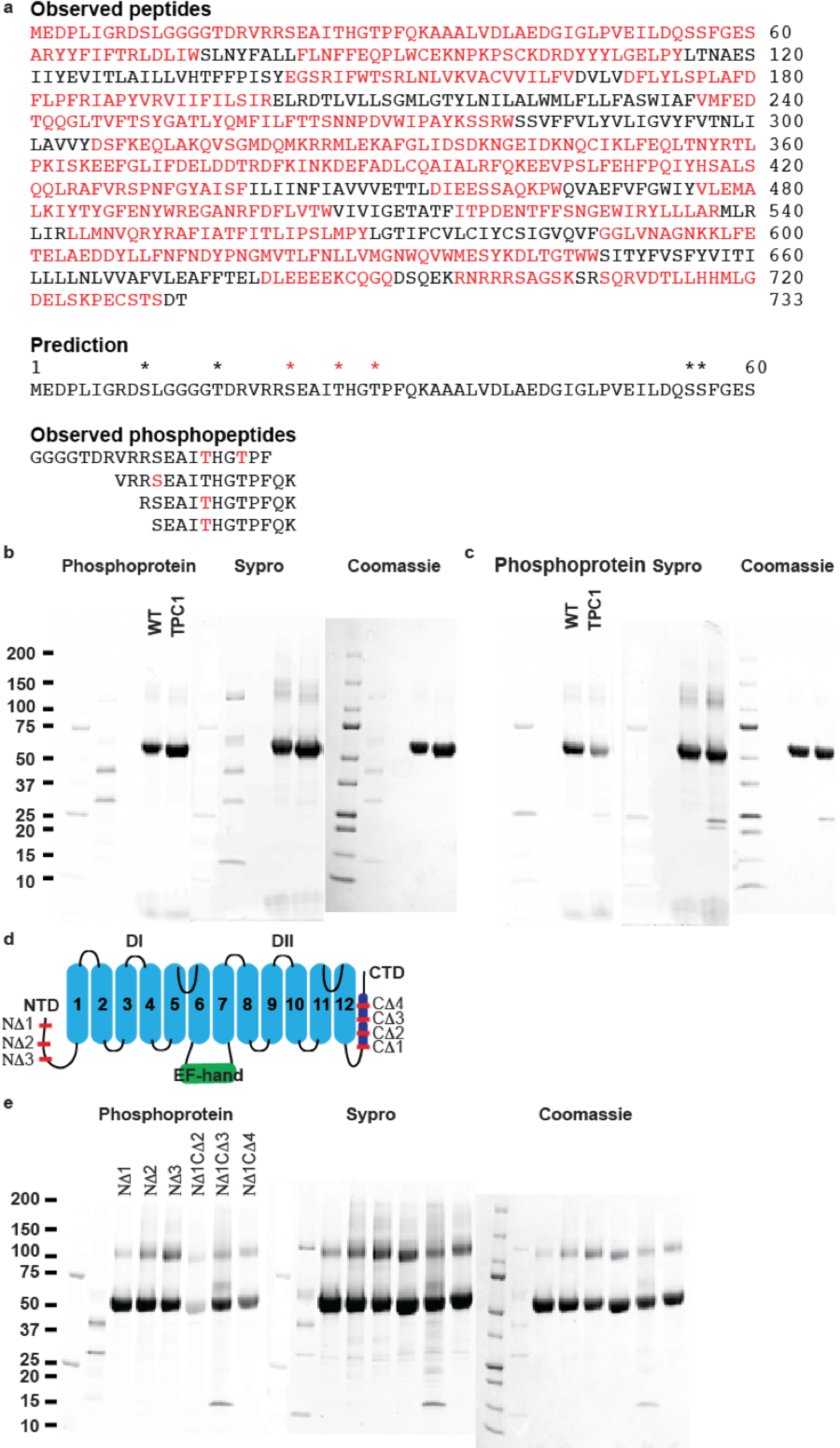
Determination of phosphorylation sites in TPC1. **a**, Electrospray ionization mass spectrometry (ESI-MS) peptide sequence coverage from an in-gel digest of wild-type (WT) TPC1 by four enzyme combinations (Trypsin/Asp-N, Trypsin/Glu-C, Lys-C, or Chymotrypsin). Measured peptides (Top, red highlight), (Middle) predicted (*) and observed (*) phosphorylation sites are shown. (Bottom) Observed phosphopeptides are listed with the sites of phosphorylation colored red. **b-e**, Polyacrylamide gels of purified TPC1 (10μg) stained with phosphoprotein-specific probe ProDiamond-Q, SyproRuby, or Coomassie (left to right). Molecular weights of standards are indicated in kDa. units to the left. First two lanes are PrecisionPlus protein molecular weight standards and PeppermintStick phosphoprotein molecular weight standards. **b**, WT TPC1 and crystallographic TPC1. **c**, Untreated and treated WT TPC1 with Lambda phosphatase for 1 hour at 25°C. **d**, Schematic of TPC1 truncations (NΔ1; 2-11, NΔ2; 2-21, NΔ3; 2-30, CΔ1; 682-733, CΔ2; 693-733, CΔ3; 709-733, CΔ4; 724-733. **e**, Analysis of TPC1 N-and C-terminal truncations for binding to ProDiamond-Q. NA1CA1 was unstable during purification.

**Extended Data Figure 4.**
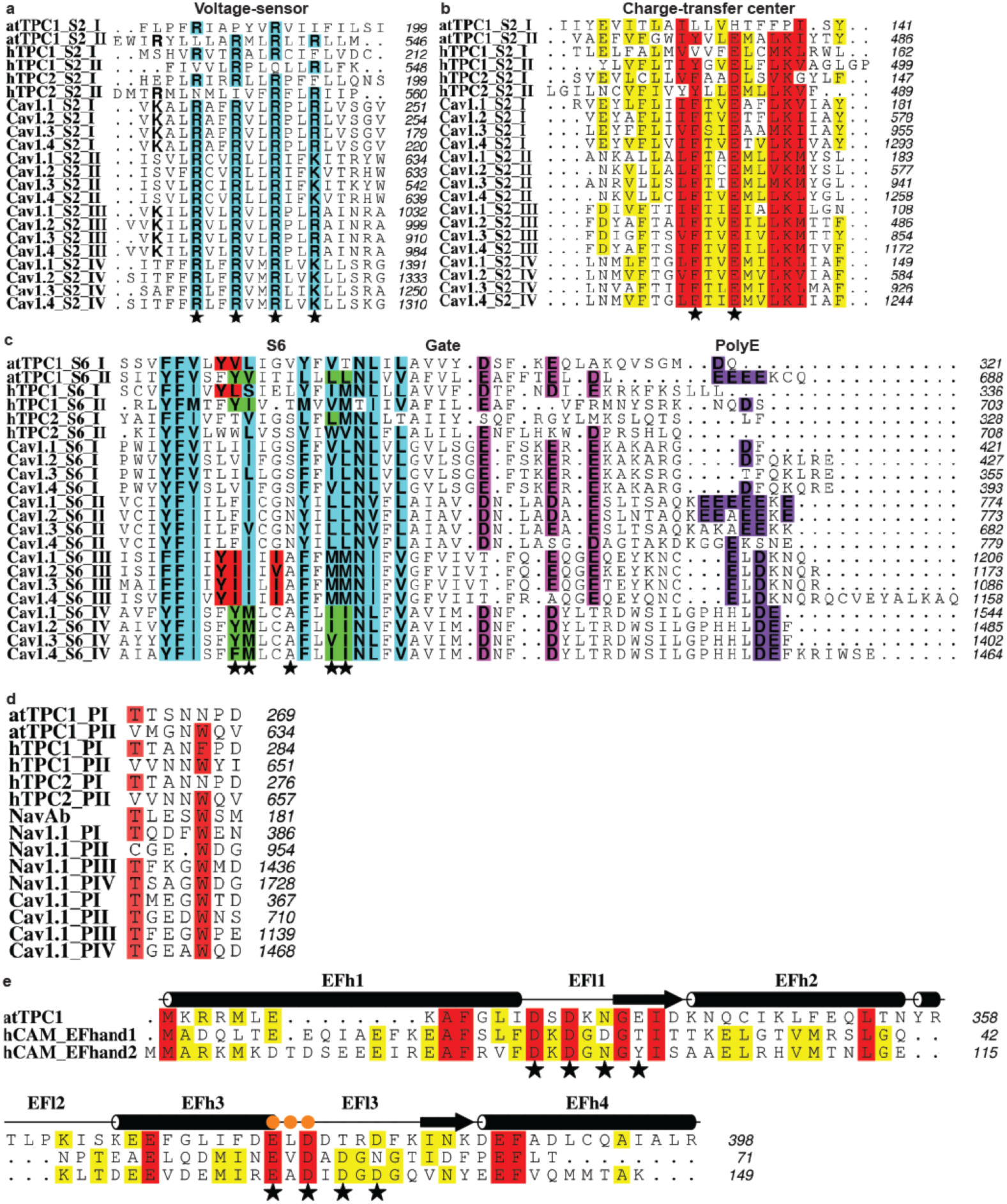
Sequence alignment of TPC1 subdomains with hTPC1, hTPC2,human Ca_v_ 1.1-1.4, Na_v_ 1.1, Na_v_Ab, and hCAM. **a**, S4 voltage-sensing segments. Conserved arginines are highlighted in cyan. Stars (⋆) mark potential voltage-sensing residues. **b**, S2segments. Stars (⋆) mark conserved charge-transfer residues. **c**, S6 segments, pore gate, and poly-E motif. Conserved hydrophobic residues are highlighted in cyan. Dihydropyridine-binding residues in the Ca_v_ S6 domain III, domain IV, and corresponding residues in TPC1 are highlighted in red and green, respectively. Stars (⋆) mark the position of phenylalkylamine drug-binding residues in Ca_v_s. Conserved residues in the gate and poly-E motif are highlighted in magenta and purple, respectively. **d**, Pore loops. Conserved residues are highlighted in red. **e**, Alignment of hCAM EF-hand domains (EF-hand 1, residues 1-71; EF-hand 2, residues 72-149)and atTPC1 (residues 322-398). Stars (⋆) mark calcium binding motifs and orange dots (•)mark the interaction site with CTD.

**Extended Data Figure 5.**
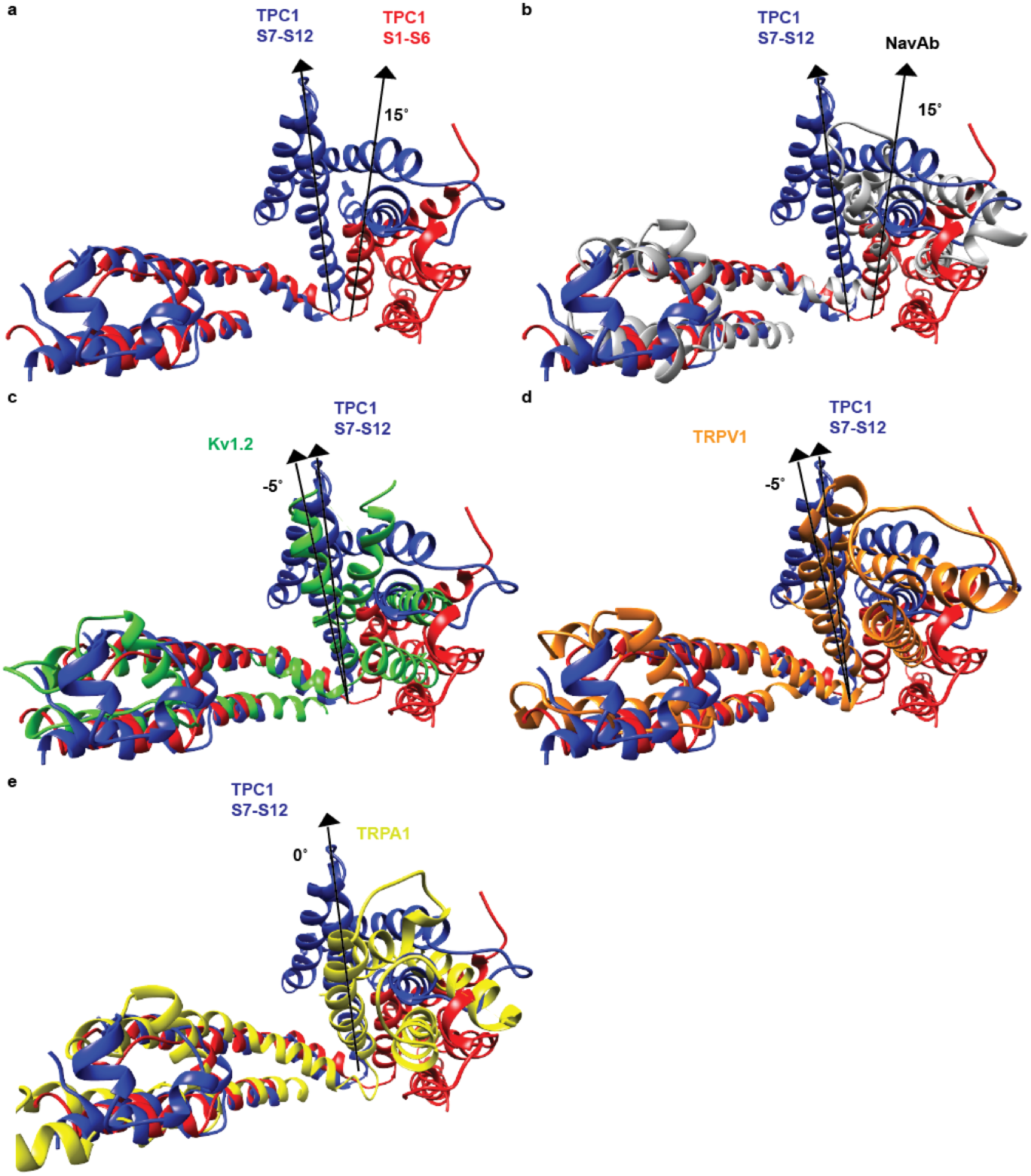
Comparison of VSDs between TPC1 and symmetrical ion channels. Structural alignment of S11-S12 segments of TPC1 domain II (blue) with S5-S6 of **a**, TPC1 domain I (red), **b**, Na_v_Ab^16^ (PDBID 3RVY, grey), **c**, K_v_1.2^15^ (PDBID 2A79, green), **d**, TRPV1^51^ (PDBID 3J5P, orange), and **e**, TRPA1^52^ (PDBID 3J9P, yellow). Angles between S4 segments with respect to S10 of TPC1 domain II are shown.

**Extended Data Figure 6.**
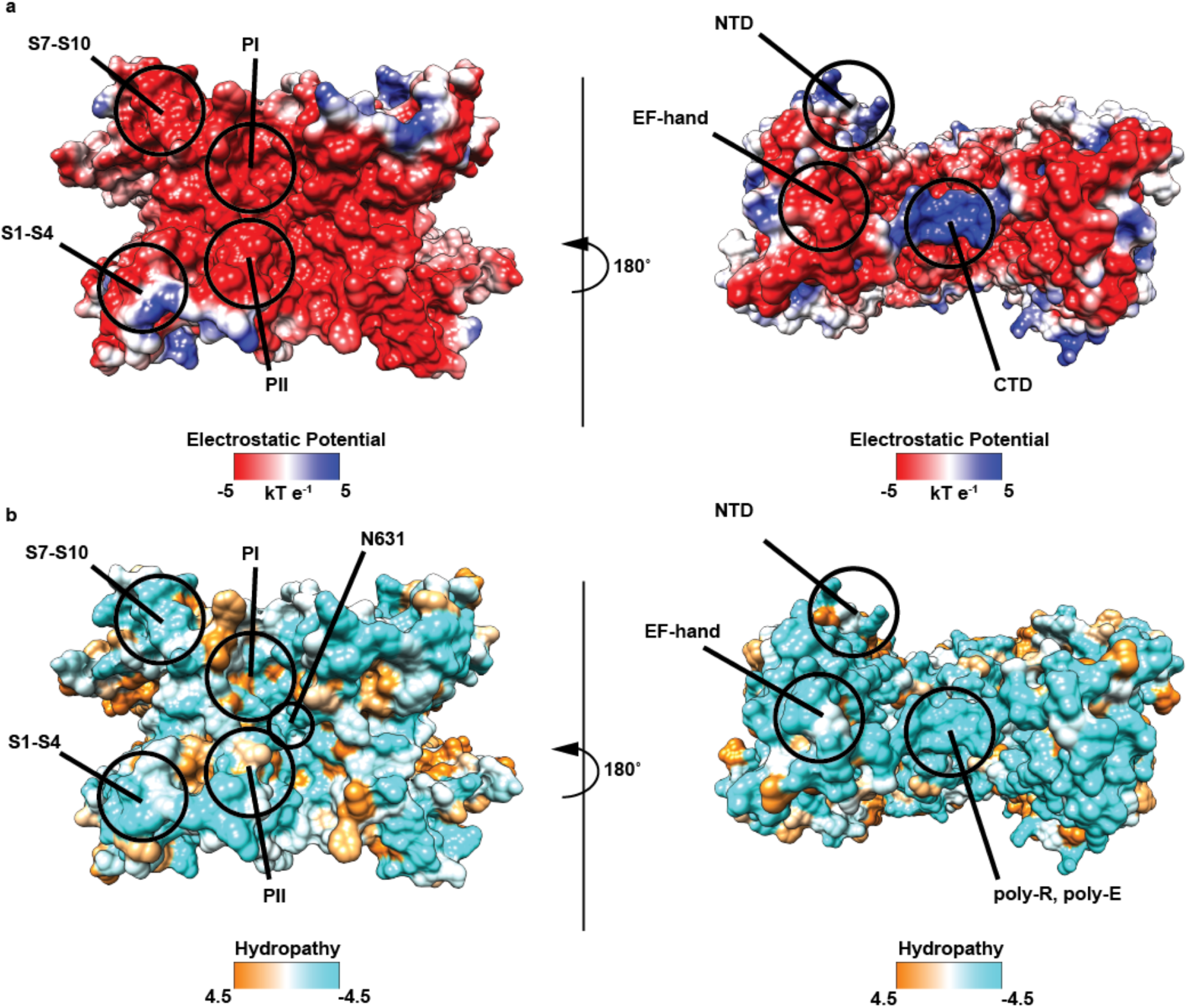
Electrostatic surfaces and Hydrophobic surfaces. (left to right) Top from the luminal side and bottom from the cytoplasmic side views of an **a**, electrostatic surface representation and **b**, surface representation colored according to Kyte-Doolittle hydropathy of TPC1. Important domains and residues are labeled. Electrostatic potential and Kyte-Doolittle hydropathy without any bound ions was generated using Chimera^46^. The EF-hand and NTD domains are negatively charged and bind cations. The poly-R motif accounts for the positively charged region in the CTD.

**Extended Data Figure 7.**
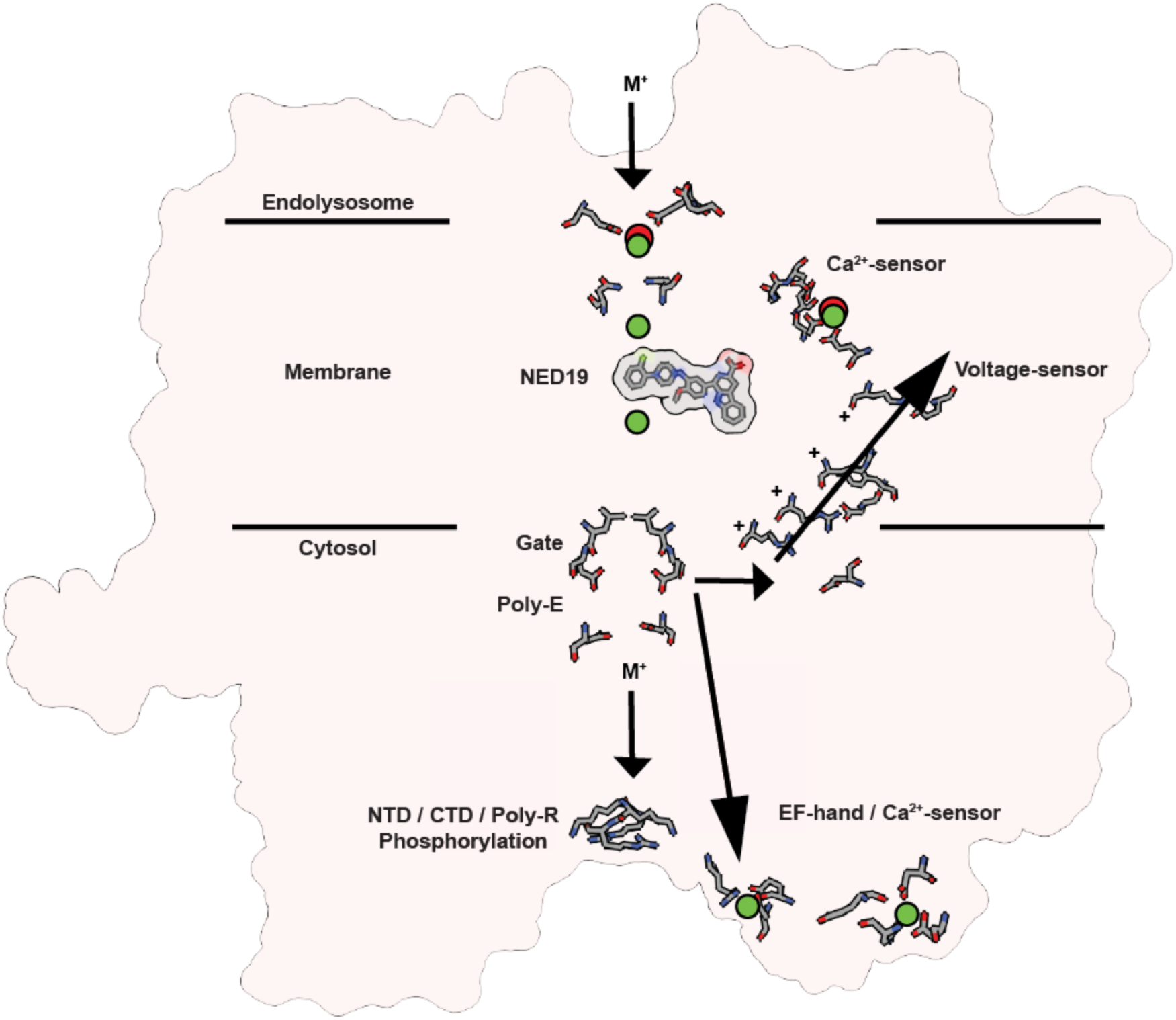
Mechanism for TPC gating. A schematic summarizing structural features of TPC1 that suggest mechanisms for voltage-sensing, ion permeation, inhibition by NED 19, lumenal Ca^2+^-inhibition, and cytosolic Ca^2+^-activation (EF-hand), and phosphoregulation (NTD/CTD). Ca^2+^ binding (green) and lanthanide (red) binding sites are shown. An ion permeation pathway through the putative selectivity filter, gate, and poly-E and poly-R motifs are summarized. **M+** represents a general cation (Na^+^, K^+^, Ca^2+^) and + signs are gating charges.

## Supplementary Information

**Structure, inhibition, and regulation of two-pore channel TPC1 from *Arabidopsis thaliana*** Alexander F. Kintzer and Robert M. Stroud

### Supplementary Discussion

#### Introduction

Two pore channels (TPCs) comprise members of the superfamily of voltage-gated ion channels^1,2^. Electrophysiological recordings from whole plant vacuoles identified a voltage-gated, slow-vacuolar (SV), calcium-activated calcium current^3^. The gene encoding the SV channel was later identified and named TPC1 in rats^4^ and plants^5^ based on sequence homology to human voltage-gated Na^+^ (Na_v_) and Ca^2+^ (Ca_v_) channels^6^. Analogous to Na_v_ and Ca_v_ channels, which encode four tandem pore-forming domains on a single chain, TPCs encode two tandem domains separated by a cytosolic domain. Plant TPC1 contains cytosolic EF-hand domains homologous to calmodulin (hCAM, ref.^7^), replaced with a domain of unknown function in human TPCs (Extended Data Figure 1). TPCs show pharmacological similarities to Na_v_ and Ca_v_ channels^8^. In plants TPC1 regulates stomatal movement, germination, and long-range stress-induced Ca^2+^ waves^9^^−^^11^. They are non-selective ion channels that conduct Ca^2+^, Na^+^, and K^+^ ions, whereas human TPC1 is Na^+^-selective and human TPC2 conducts Ca^2+^, Na^+^, and H^+^ ions^12^. TPC1 functions as the major plant vacuolar calcium-induced calcium release channel^13^. In animals, however, Ca^2+^-release by TPC2 from the endolysosome, is subsequently amplified by ryanodine and inositol triphosphate receptors embedded in the endoplasmic reticulum^14^. Cytosolic Ca^2+^ ion concentration uniquely potentiates plant TPCs mediated by distinct calmodulin-like domains^7^, not present in mammalian TPCs^4^. TPCs dimerize to form quasi-tetrameric channels composed of alternating copies of the two divergent tandem pore-forming domains (D1 and D2)^15^. They have two homologous Shaker-like voltage-sensing domains in transmembrane segments S1-S4, and S7-S10, pore loops in S5-S6 and S11-S12, and activation gates following S6 and S12. D1 and D2 share higher sequence identity with particular domains in Ca_v_ and Na_v_ channels, than with one another^6^, suggesting that they are evolutionarily derived from a common ancestor and may share functional mechanisms.

TPCs sense and respond to numerous luminal and cytosolic agents that regulate ion release from the vacuole, or the endolysosome^3,12,16-^24^^. These include Ca^2+^, pH, and cellular redox potential. Plant TPC1 is inhibited by polyunsaturated fatty acids^20^, whereas the lysosome-specific second messenger lipid PI(3,5)P_2_ increases the open probability of human TPC2, but not TPC1 channels^21^.

The N-terminal domain of plant TPC1 contains a vacuolar localization signal sequence that is conserved with the endolysosomal targeting signal sequence in animals^25,26^. A binding site on the N-terminus of human TPCs for Rab GTPases serves to regulate trafficking^27^. The C-terminus may form a hub for binding of apoptotic machinery^28^. However, it is not known yet whether plant TPCs participate in protein-protein interactions via the N-or C-termini.

Both plant and human TPCs are known to respond to phosphoregulation^29^^−^^32^. CAM-like dependent kinase, for instance, activates plant TPC1, but phosphorylation at a distinct site by a PKA, PKC, or PKG-like kinase inhibits the channel. mTOR kinase^33^ phosphorylates human TPCs and so tunes the release of nutrients and ions from the endolysosome in response to cellular growth signals, stress, or starvation^30^. Human diseases, including diabetes, obesity, cancer, and age-related diseases result from defective signaling both upstream and downstream of mTOR^33^. Targeting the TPC-mTOR complex is therefore an avenue for treatment of various human diseases. Hence phosphoregulation plays an important role in TPC function *in vivo*, as with other ion channels.

Human TPCs form a stable complex with mTOR kinase, facilitating direct phosphorylation of the channel and stabilizing the closed state^30^. MAPK, JNK, and P38 kinases^34,35^ also inhibit the conductance of hTPC2 by phosphorylation^29^. LRRK2 kinase has been shown to bind and regulate hTPC2^36^, though effects on the channel conductance have not been measured. NTDh2 in hTPC2 contains a Rab GTPase-binding motif, not conserved in hTPC1, that facilitates interaction with Rabs and regulates TPC1 trafficking and channel function^27^. The interaction of plant TPCs with Rab GTPases has not been tested, but it is conceivable that binding to NTDh2 could affect channel opening. Thus the NTD plays important roles in channel function, phosphoregulation, and intracellular trafficking.

In most eukaryotic cells, though apparently not in plants or yeast, nicotinic acid adenine dinucleotide phosphate (NAADP) triggers the release of Ca^2+^ ions from acidic intracellular stores, causing temporary increases in the cytosolic Ca^2+^ ion concentration^37,38^. Ryanodine receptors further amplify these Ca^2+^ transients to produce changes in important biological processes ranging from cell differentiation to cardiac function^32,39,40^. NAADP triggers opening of hTPC2 channels indirectly^21,22^, probably via a complex with an as yet unidentified NAADP-binding protein^27^.

Filoviruses, such as Ebola virus require host-cell receptors, endocytosis, proteolytic cleavage, and fusion with the endolysosomal membrane for release of viral material into the cytoplasm^23,41−^^43^. TPC antagonists disrupt the intracellular release of Na^+^ or Ca^2+^ ions from the endolysosome and inhibit the trafficking of Filoviruses, preventing infection^23^. Antagonists trans-NED19 (NED19; ref.^44^), and approved medications of the benzothiazepine, dihydropyridine (DHP), and phenylalkylamine (PAA) classes of L-type Ca_v_ antagonists inhibit TPCs directly by blocking channel opening^22^. The most potent anti-viral TPC antagonist, tetraandrine, provides an effective treatment for Ebola disease in animals^23^. Both DHP and PAA pharmacophores inhibit plant TPC channels, owing to conserved hydrophobic residues in transmembrane segment S6^45^^−^^47^. Thus, understanding the basis for inhibition of plant and human TPCs by pharmacophores may help determine the molecular underpinnings and novel avenues for the treatment of critical human diseases.

#### Determination of Phosphorylation sites in TPC1

We determined that purified plant TPC1 is phosphorylated by binding of a phosphoprotein-specific probe (ProDiamond-Q) to both wild-type TPC1 and TPC1 (Extended Data Figure 3). Treatment with a phosphatase reduces binding of the probe, indicating that the sites of phosphorylation are mostly solvent accessible and reversible (Extended Data Figure 3b).

We determined sites of phosphorylation in plant TPC1 using a combination of mass spectrometry and binding of ProDiamond-Q to N-and C-terminal truncations. Electrospray-ionization mass spectrometry (ESI-MS) analysis of peptides derived from proteolytic digests of gel bands (trypsin/Asp-N, Chymotrypsin, and Lys-C) identified peptides for 87.2% of residues in non-transmembrane regions, 38.1% in transmembrane regions, and 72.6% overall (Extended Data Figure 3a). The ESI-MS data covers most of the predicted solvent exposed sites of phosphorylation, with the exception of several sites in the NTD, P1, S7-S8, CTD, and transmembrane segments. To detect phosphorylation sites within the gaps in ESI-MS coverage, N-(ΔN1-3) and C-terminal (ΔC1-4) truncations of TPC1 (Extended Data Figure 3e) were analyzed for binding to ProDiamond-Q. Deleting part of the C-terminus (ΔN1ΔC2) abrogated binding of the probe, suggesting that the CTD may contain sites of phosphorylation and localizes the likely sites to a patch of serine residues in the CTD.

#### Modulation of VSDs by ions

Voltage-gated ion channels present charged residues in S2 and S4 that respond to the transmembrane electric field. Movement of S4 is mechanically coupled to gating of the channel, requiring at least three regularly spaced arginine residues in S4, and a counter anion in S2^48,49^. VSDs 1 and 2 in plant TPC1 share roughly 25% sequence identity with each other and with the VSDs in human TPCs and Ca_v_s (Extended Data Fig. 4).

#### Ion selectivity and pore blockade

Lanthanide ions block plant TPCs^10,16^. The Yb^3+^-bound TPC_cryst_ structure shows direct blockade of the upper pore loop 2 coordinated by four carboxyls (Fig. 5, Extended Data Figure 2c), suggesting that lanthanides may block TPCs by binding near the pore mouth.

#### Cytoplasmic sensory domains

The plant vacuole functions as the primary intracellular Ca^2+^ store and comprises 90% of the cell volume^3,5^. Plant TPCs coordinate Ca^2+^-activated release of Ca^2+^ ions from the vacuole to the cytoplasm. The CTD plays a critical role in plant TPCs^26^, though least conserved across species (Extended Data Fig. 1). Deletion of the CTD results in inactive plant TPC1 channels. Plant TPC channels are known to be sensitive to cytoplasmic redox potential^19^. A Hg^2+^ site (Hg1) binds to a cysteine (C347) in EFh1. This site could reasonably function as the redox sensor, which facilitates channel opening in response to lowering of cytoplasmic pH and more reducing conditions. On the luminal side of the membrane a CxxCxxC (C574, C577, C580) motif in S11 (Extended Data Fig. 1) is characteristic of an iron-sulfur cluster-binding site and often linked to redox processes in the endoplasmic reticulum. The function of this site in plant biology has not been described. And the CxxCxxC motif is not conserved in human TPCs.

**Supplementary Table 1.**
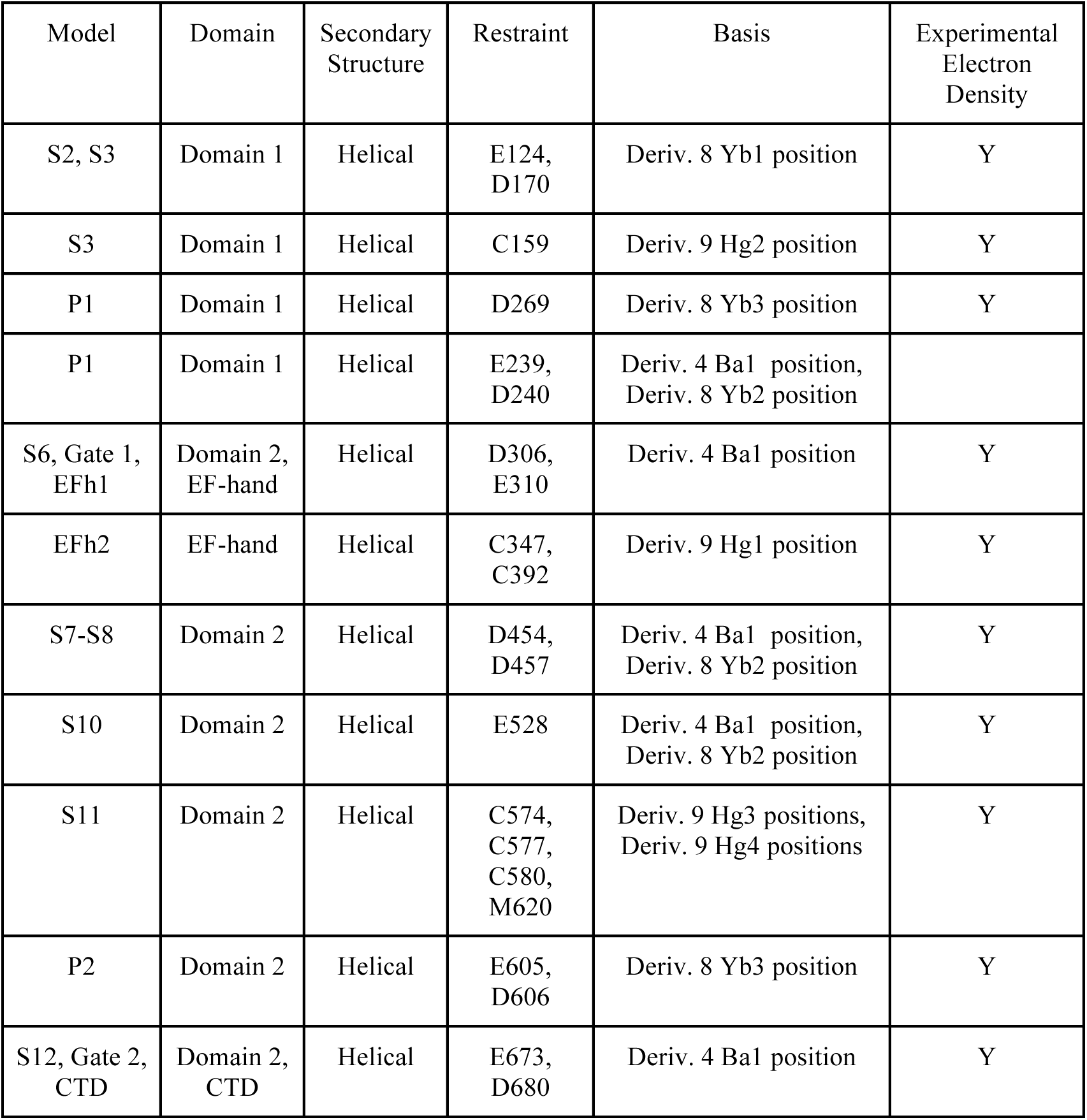
Building restraints for TPC1. Experimental restraints used for objective assignment of the TPC1 sequence to electron density.

## References

1. Hedrich, R. & Neher, E. Cytoplasmic calcium regulates voltage-dependent ion channels in plant vacuoles. Nature 329, 833–836 (1987).

2. Calcraft, P. J. etal. NAADP mobilizes calcium from acidic organelles through two-pore channels. Nature 459, 596–600 (2009).

3. Hedrich, R. & Marten, I. TPC1 - SV Channels Gain Shape. Mol. Plant 4, 428–441 (2011).

4. Patel, S. Function and dysfunction of two-pore channels. Sci Signal 8, re7–re7 (2015).

5. Wang, X. etal. TPC proteins are phosphoinositide-activated sodium-selective ion channels in endosomes and lysosomes. Cell 151, 372–383 (2012).

6. Pitt, S. J. etal. TPC2 Is a Novel NAADP-sensitive Ca2+ Release Channel, Operating as a Dual Sensor of Luminal pH and Ca2+ J. Biol. Chem. 285, 35039–35046 (2010).

7. Sakurai, Y. et al. Two-pore channels control Ebola virus host cell entry and are drug targets for disease treatment. Science 347, 995–998 (2015).

8. Hedrich, R., Flügge, U. I. & Fernandez, J. M. Patch-clamp studies of ion transport in isolated plant vacuoles. FEBS Lett. 204, 228–232 (1986).

9. Cang, C., Bekele, B. & Ren, D. The voltage-gated sodium channel TPC1 confers endolysosomal excitability. Nat. Chem. Biol. 10, 463–469 (2014).

10. Guo, J. et al. Structure of the voltage-gated two-pore channel TPC1 from Arabidopsis thaliana. Nature advance online publication, (2015).

11. Beyhl, D. etal. The fou2 mutation in the major vacuolar cation channel TPC1 confers tolerance to inhibitory luminal calcium. Plant J. 58, 715–723 (2009).

12. Cang, C. et al. mTOR Regulates Lysosomal ATP-Sensitive Two-Pore Na+ Channels to Adapt to Metabolic State. Cell 152, 778–790 (2013).

13. Morgan, A. J., Davis, L. C., Ruas, M. & Galione, A. TPC: the NAADP discovery channel? Biochem. Soc. Trans. 43, 384–389 (2015).

14. Schulze, C., Sticht, H., Meyerhoff, P. & Dietrich, P. Differential contribution of EF-hands to the Ca2+-dependent activation in the plant two-pore channel TPC1. Plant J. 68, 424–432 (2011).

15. Long, S. B., Campbell, E. B. & Mackinnon, R. Crystal structure of a mammalian voltage-dependent Shaker family K+ channel. Science 309, 897–903 (2005).

16. Payandeh, J., Scheuer, T., Zheng, N. & Catterall, W. A. The crystal structure of a voltage-gated sodium channel. Nature 475, 353–358 (2011).

17. Naylor, E. etal. Identification of a chemical probe for NAADP by virtual screening. Nat. Chem. Biol. 5, 220–226 (2009).

18. Kamb, A., Iverson, L. E. & Tanouye, M. A. Molecular characterization of Shaker, a Drosophila gene that encodes a potassium channel. Cell 50, 405–413 (1987).

19. Papazian, D. M., Schwarz, T. L., Tempel, B. L., Jan, Y. N. & Jan, L. Y. Cloning of genomic and complementary DNA from Shaker, a putative potassium channel gene from Drosophila. Science 237, 749–753 (1987).

20. Wu, J. et al. Structure of the voltage-gated calcium channel Cav1.1 complex. Science 350, aad2395 (2015).

21. Catterall, W. A., Perez-Reyes, E., Snutch, T. P. & Striessnig, J. International Union of Pharmacology. XLVIII. Nomenclature and Structure-Function Relationships of Voltage-Gated Calcium Channels. Pharmacol. Rev. 57, 411–425 (2005).

22. Brailoiu, E. et al. Essential requirement for two-pore channel 1 in NAADP-mediated calcium signaling. J. Cell Biol. 186, 201–209 (2009).

23. Peterson, B. Z., Tanada, T. N. & Catterall, W. A. Molecular Determinants of High Affinity Dihydropyridine Binding in L-type Calcium Channels. J. Biol. Chem. 271, 5293–5296 (1996).

24. Pitt, S. J., Lam, A. K. M., Rietdorf, K., Galione, A. & Sitsapesan, R. Reconstituted human TPC1 is a proton-permeable ion channel and is activated by NAADP or Ca2+. Sci. Signal. 7, ra46(2014).

25. Tang, L. et al. Structural basis for Ca2+ selectivity of a voltage-gated calcium channel. Nature 505, 56–61 (2014).

26. Armstrong, C. M. Time Course of TEA+-Induced Anomalous Rectification in Squid Giant Axons. J. Gen. Physiol. 50, 491–503 (1966).

27. Holmgren, M., Smith, P. L. & Yellen, G. Trapping of Organic Blockers by Closing of Voltage-dependent K+ Channels Evidence for a Trap Door Mechanism of Activation Gating. J. Gen. Physiol. 109, 527–535 (1997).

28. Perozo, E., Cortes, D. M. & Cuello, L. G. Three-dimensional architecture and gating mechanism of a K+ channel studied by EPR spectroscopy. Nat. Struct. Mol. Biol. 5, 459–469 (1998).

29. Clayton, G. M., Altieri, S., Heginbotham, L., Unger, V. M. & Morais-Cabral, J. H. Structure of the transmembrane regions of a bacterial cyclic nucleotide-regulated channel. Proc. Natl. Acad. Sci. 105, 1511–1515 (2008).

30. Zhou, Y., Morais-Cabral, J. H., Kaufman, A. & MacKinnon, R. Chemistry of ion coordination and hydration revealed by a K+ channel-Fab complex at 2.0 | [angst] | resolution. Nature 414, 43–48 (2001).

31. Mumberg, D., Müller, R. & Funk, M. Regulatable promoters of Saccharomyces cerevisiae: comparison of transcriptional activity and their use for heterologous expression. Nucleic Acids Res. 22, 5767–5768 (1994).

32. Hattori, M., Hibbs, R. E. & Gouaux, E. A Fluorescence-Detection Size-Exclusion Chromatography-Based Thermostability Assay for Membrane Protein Precrystallization Screening. Structure 20, 1293–1299 (2012).

33. Gourdon, P. et al. HiLiDe—Systematic Approach to Membrane Protein Crystallization in Lipid and Detergent. Cryst. Growth Des. 11, 2098–2106 (2011).

34. Winn, M. D. et al. Overview of the CCP4 suite and current developments. Acta Crystallogr. D Biol. Crystallogr. 67, 235–242 (2011).

35. Strong, M. etal. Toward the structural genomics of complexes: Crystal structure of a PE/PPE protein complex from Mycobacterium tuberculosis. Proc. Natl. Acad. Sci. 103, 8060–8065 (2006).

36. Kabsch, W. Automatic processing of rotation diffraction data from crystals of initially unknown symmetry and cell constants. J. Appl. Crystallogr. 26, 795–800 (1993).

37. Sheldrick, G. M. Experimental phasing with SHELXC/D/E: combining chain tracing with density modification. Acta Crystallogr. D Biol. Crystallogr. 66, 479–485 (2010).

38. Bricogne, G., Vonrhein, C., Flensburg, C., Schiltz, M. & Paciorek, W. Generation, representation and flow of phase information in structure determination: recent developments in and around SHARP 2.0. Acta Crystallogr. D Biol. Crystallogr. 59, 2023–2030 (2003).

39. Abrahams, J. P. & Leslie, A. G. W. Methods used in the structure determination of bovine mitochondrial F1 ATPase. Acta Crystallogr. D Biol. Crystallogr. 52, 30–42 (1996).

40. Cowtan, K. dm: An automated procedure for phase improvement by density modification. Jt. CCP4 ESF-EACBMNewsl. Protein Crystallogr. 31, 34–38 (1994).

41. Pedersen, B. P., Morth, J. P. & Nissen, P. Structure determination using poorly diffracting membrane-protein crystals: the H+-ATPase and Na+,K+-ATPase case history. Acta Crystallogr. D Biol. Crystallogr. 66, 309–313 (2010).

42. Emsley, P. & Cowtan, K. Coot: model-building tools for molecular graphics. Acta Crystallogr. D Biol. Crystallogr. 60, 2126–2132 (2004).

43. Adams, P. D. et al. PHENIX: a comprehensive Python-based system for macromolecular structure solution. Acta Crystallogr. D Biol. Crystallogr. 66, 213–221 (2010).

44. Edgar, R. C. MUSCLE: multiple sequence alignment with high accuracy and high throughput. Nucleic Acids Res. 32, 1792–1797 (2004).

45. Bond, C. S. & Schüttelkopf, A. W. ALINE: a WYSIWYG protein-sequence alignment editor for publication-quality alignments. Acta Crystallogr. D Biol. Crystallogr. 65, 510–512 (2009).

46. Pettersen, E. F. et al. UCSF Chimera--a visualization system for exploratory research and analysis. J. Comput. Chem. 25, 1605–1612 (2004).

47. Smart, O. S., Neduvelil, J. G., Wang, X., Wallace, B. A. & Sansom, M. S. P. HOLE: A program for the analysis of the pore dimensions of ion channel structural models. J. Mol. Graph. 14, 354–360 (1996).

48. Jones, D. T. Protein secondary structure prediction based on position-specific scoring matrices 1. J. Mol. Biol. 292, 195–202 (1999).

49. Cuff, J. A., Clamp, M. E., Siddiqui, A. S., Finlay, M. & Barton, G. J. JPred: a consensus secondary structure prediction server. Bioinformatics 14, 892–893 (1998).

50. Blom, N., Sicheritz-Pontén, T., Gupta, R., Gammeltoft, S. & Brunak, S. Prediction of post-translational glycosylation and phosphorylation of proteins from the amino acid sequence. PROTEOMICS 4, 1633–1649 (2004).

51. Liao, M., Cao, E., Julius, D. & Cheng, Y. Structure of the TRPV1 ion channel determined by electron cryo-microscopy. Nature 504, 107–>112 (2013).

52. Paulsen, C. E., Armache, J.-P., Gao, Y., Cheng, Y. & Julius, D. Structure oftheTRPA1ion channel suggests regulatory mechanisms. Nature 520, 511–517 (2015).

## Supplementary Information References

1. Catterall, W. A. Structure and Function of VoltageAGated Ion Channels. Annu.%Rev.% Biochem. 64, 493–531 (1995).

2. Yu, F. H. & Catterall, W. A. The VGLAChanome: A Protein Superfamily Specialized for Electrical Signaling and Ionic Homeostasis. Sci%STKE 2004, re15 (2004).

3. Hedrich, R. & Neher, E. Cytoplasmic calcium regulates voltageAdependent ion channels in plant vacuoles. Nature 329, 833–836 (1987).

4. Ishibashi, K., Suzuki, M. & Imai, M. Molecular Cloning of a Novel Form (TwoARepeat) Protein Related to VoltageAGated Sodium and Calcium Channels. Biochem.%Biophys.%Res.% Commun. 270, 370–376 (2000).

5. Furuichi, T., Cunningham, K. W. & Muto, S. A Putative Two Pore Channel AtTPC1 Mediates Ca2+ Flux in Arabidopsis Leaf Cells. Plant%Cell%Physiol. 42, 900–905 (2001).

6. Rahman, T. et%al. TwoApore channels provide insight into the evolution of voltageAgated Ca2+ and Na+ channels. Sci%Signal 7, ra109–ra109 (2014).

7. Schulze, C., Sticht, H., Meyerhoff, P. & Dietrich, P. Differential contribution of EFAhands to the Ca2+Adependent activation in the plant twoApore channel TPC1. Plant%J. 68, 424–432 (2011).

8. Hashimoto, K., Saito, M., Matsuoka, H., Iida, K. & Iida, H. Functional Analysis of a Rice Putative VoltageADependent Ca2+ Channel, OsTPC1, Expressed in Yeast Cells Lacking its Homologous Gene CCH1. Plant Cell Physiol. 45, 496–500 (2004).

9. Peiter, E. etal. The vacuolar Ca2+Aactivated channel TPC1 regulates germination and stomatal movement. Nature 434, 404–408 (2005).

10. Choi, W.AG., Toyota, M., Kim, S.AH., Hilleary, R. & Gilroy, S. Salt stressAinduced Ca2+ waves are associated with rapid, longAdistance rootAtoAshoot signaling in plants. Proc. Natl. Acad. Sci. 111, 6497–6502 (2014).

11. Gilroy, S. etal. A tidal wave of signals: calcium and ROS at the forefront of rapid systemic signaling. Trends Plant Sci. 19, 623–630 (2014).

12. Pitt, S. J. et al. TPC2 Is a Novel NAADPAsensitive Ca2+ Release Channel, Operating as a Dual Sensor of Luminal pH and Ca2+. J. Biol. Chem. 285, 35039–35046 (2010).

13. Bewell, M. A., Maathuis, F. J. M., Allen, G. J. & Sanders, D. CalciumAinduced calcium release mediated by a voltageAactivated cation channel in vacuolar vesicles from red beet. FEBSLett. 458, 41–44 (1999).

14. Gerasimenko, J. V. et al. Both RyRs and TPCs are required for NAADPAinduced intracellular Ca2+ release. Cell Calcium doi:10.1016/j.ceca.2015.05.005

15. Rietdorf, K. etal. TwoApore Channels Form HomoA and Heterodimers. J. Biol. Chem. 286, 37058–37062(2011).

16. Kawano, T. et al. Aluminum as a specific inhibitor of plant TPC1 Ca2+ channels. Biochem. Biophys. Res. Commun. 324, 40–45 (2004).

17. Beyhl, D. et al. The fou2 mutation in the major vacuolar cation channel TPC1 confers tolerance to inhibitory luminal calcium. Plant%J. 58, 715–723 (2009).

18. Carpaneto, A., Cantù, A. M. & Gambale, F. Redox agents regulate ion channel activity in vacuoles from higher plant cells. FEBS Lett. 442, 129–132 (1999).

19. ScholzAStarke, J. et al. RedoxAdependent modulation of the carrot SV channel by cytosolic pH. FEBS Lett. 576, 449–454 (2004).

20. Gutla, P. V. K., Boccaccio, A., De Angeli, A., Gambale, F. & Carpaneto, A. Modulation of plant TPC channels by polyunsaturated fatty acids. J. Exp. Bot. 63, 6187–6197 (2012).

21. Wang, X. etal. TPC proteins are phosphoinositideA activated sodiumAselective ion channels in endosomes and lysosomes. Cell 151, 372–383 (2012).

22. Ruas, M. et al. Expression of Ca2+Apermeable twoApore channels rescues NAADP signalling in TPCAdeficient cells. EMBOJ. 34, 1743–1758 (2015).

23. Sakurai, Y. et al. TwoApore channels control Ebola virus host cell entry and are drug targets for disease treatment. Science 347, 995–998 (2015).

24. Genazzani, A. A. etal. Pharmacological properties of the Ca2+Arelease mechanism sensitive to NAADP in the sea urchin egg. Br. J. Pharmacol. 121, 1489–1495 (1997).

25. Churamani, D., Hooper, R., Rahman, T., Brailoiu, E. & Patel, S. The NAterminal region of twoApore channel 1 regulates trafficking and activation by NAADP. Biochem.J. 453, 147–151(2013).

26. Larisch, N., Schulze, C., Galione, A. & Dietrich, P. An NATerminal Dileucine Motif Directs TwoAPore Channels to the Tonoplast of Plant Cells. Traffic 13, 1012–1022 (2012).

27. LinAMoshier, Y. et al. The TwoApore channel (TPC) interactome unmasks isoformA specific roles for TPCs in endolysosomal morphology and cell pigmentation. Proc. Natl. Acad. Sci. 111, 13087–13092 (2014).

28. Lam, A. K. M., Galione, A., Lai, F. A. & Zissimopoulos, S. HaxA1 identified as a twoApore channel (TPC)Abinding protein. FEBS Lett. 587, 3782–3786 (2013).

29. Jha, A., Ahuja, M., Patel, S., Brailoiu, E. & Muallem, S. Convergent regulation of the lysosomal twoApore channelA2 by Mg2+, NAADP, PI(3,5)P2 and multiple protein kinases. EMBOJ. 33, 501–511 (2014).

30. Cang, C. et al. mTOR Regulates Lysosomal ATPASensitive TwoAPore Na+ Channels to Adapt to Metabolic State. Cell 152, 778–790 (2013).

31. Bethke, P. C. & Jones, R. L. Reversible protein phosphorylation regulates the activity of the slowAvacuolar ion channel. Plant J. 11, 1227–1235 (1997).

32. Morgan, A. J., Davis, L. C., Ruas, M. & Galione, A. TPC: the NAADP discovery channel? Biochem. Soc. Trans. 43, 384–389 (2015).

33. Laplante, M. & Sabatini, D. M. mTOR Signaling in Growth Control and Disease. Cell 149, 274–293(2012).

34. Wullschleger, S., Loewith, R. & Hall, M. N. TOR Signaling in Growth and Metabolism. Cell 124, 471–484 (2006).

35. Johnson, G. L. & Lapadat, R. MitogenAActivated Protein Kinase Pathways Mediated by ERK, JNK, and p38 Protein Kinases. Science 298, 1911–1912 (2002).

36. GómezASuaga, P. et al. LeucineArich repeat kinase 2 regulates autophagy through a calciumAdependent pathway involving NAADP. Hum. Mol. Genet. 21, 511–525 (2012).

37. Cancela, J. M., Churchill, G. C. & Galione, A. Coordination of agonistAinduced Ca2+A signalling patterns by NAADP in pancreatic acinar cells. Nature 398, 74–76 (1999).

38. Calcraft, P. J. et al. NAADP mobilizes calcium from acidic organelles through twoA pore channels. Nature 459, 596–600 (2009).

39. Galione, A. A primer of Ca2+ signalling: From sea urchin eggs to mammalian cells. Cell Calcium 58, 27–47 (2015).

40. Patel, S. Function and dysfunction of twoApore channels. Sci Signal 8, re7–re7 (2015).

41. Lee, J. E. & Saphire, E. O. Ebolavirus glycoprotein structure and mechanism of entry. Future Virol. 4, 621–635 (2009).

42. Carette, J. E. et al. Ebola virus entry requires the cholesterol transporter NiemannA Pick C1. Nature 477, 340–343 (2011).

43. Chandran, K., Sullivan, N. J., Felbor, U., Whelan, S. P. & Cunningham, J. M. Endosomal Proteolysis of the Ebola Virus Glycoprotein Is Necessary for Infection. Science 308, 1643–1645 (2005).

44. Naylor, E. et al. Identification of a chemical probe for NAADP by virtual screening. Nat. Chem. Biol. 5, 220–226 (2009).

45. Hockerman, G. H., Johnson, B. D., Abbott, M. R., Scheuer, T. & Catterall, W. A. Molecular Determinants of High Affinity Phenylalkylamine Block of lAtype Calcium Channels in Transmembrane Segment IIIS6 and the Pore Region of the αSubunit. J. Biol. Chem. 272, 18759–18765 (1997).

46. Peterson, B. Z., Tanada, T. N. & Catterall, W. A. Molecular Determinants of High Affinity Dihydropyridine Binding in LAtype Calcium Channels. J. Biol. Chem. 271, 5293–5296 (1996).

47. White, P. J. Calcium channels in higher plants. Biochim. Biophys. Acta BBA I Biomembr. 1465, 171–189 (2000).

48. Seoh, S.AA., Sigg, D., Papazian, D. M. & Bezanilla, F. VoltageASensing Residues in the S2 and S4 Segments of the Shaker K+ Channel. Neuron 16, 1159–1167 (1996).

49. Aggarwal, S. K. & MacKinnon, R. Contribution of the S4 Segment to Gating Charge in the Shaker K+ Channel. Neuron 16, 1169–1177 (1996).

